# Epistasis shapes the fitness landscape of an allosteric specificity switch

**DOI:** 10.1101/2020.10.21.348920

**Authors:** Kyle K. Nishikawa, Nicholas Hoppe, Robert Smith, Craig Bingman, Srivatsan Raman

## Abstract

Epistasis is a major determinant in the emergence of novel protein function. In allosteric proteins, direct interactions between inducer-binding mutations propagate through the allosteric network, manifesting as epistasis at the level of biological function. Elucidating this relationship between local interactions and their global effects is essential to understanding evolution of allosteric proteins. We integrate computational design, structural and biophysical analysis to characterize the emergence of novel inducer specificity in an allosteric transcription factor. Adaptive landscapes of different inducers of the engineered mutant show that a few strong epistatic interactions constrain the number of viable sequence pathways, revealing ridges in the fitness landscape leading to new specificity. Crystallographic evidence shows a single mutation drives specificity by reshaping the binding pocket. Comparison of biophysical and functional landscapes emphasizes the nonlinear relationship between local inducer affinity and global function (allostery). Our results highlight the functional and evolutionary complexity of allosteric proteins.

## Introduction

Interactions between mutations direct the evolution of protein function^1^. As proteins evolve, they follow paths through the fitness landscape to reach a fitness peak that represents a novel function^2^. For N mutations required to confer novel function, there are N! possible pathways connecting start and end states. However, some pathways may not be evolutionarily favorable due to epistasis -- a phenomenon that occurs when the sequence background into which a mutation is introduced changes the functional effect of that mutation. The non-additivity due to epistasis strongly influences the sequence trajectory a protein takes to gain new function^1,3–5^. Therefore, understanding the nature of epistatic interactions is the foundation for investigating the mechanisms leading to novel protein function.

Epistasis is generally categorized as specific or nonspecific based on cause-effect relationships between the interactions of mutations and their outcome. Specific epistasis occurs between a limited number of residues that typically physically interact, leading to nonadditive changes in thermodynamically-driven biophysical properties such as protein stability or affinity^6^. Specific epistasis has been extensively investigated in protein-protein, protein-ligand, protein-DNA interactions^5,7–15^. Nonspecific epistasis is caused by mutations interacting nonadditively that affect biological function of the protein^16–19^. Such mutations can be spatially distant such as a global suppressor that can interact with many destabilizing mutations with low pairing specificity^4,20,21^.

In this study, we examine the role of epistasis in the evolution of function in an allosteric protein. Allostery is a fundamental mechanism by which proteins recognize environmental cues (such as binding of an inducer or effector) within a localized region resulting in modulation of function at a distal site^22,23^. Mutations in the binding pocket that trigger the allosteric network have the potential to create new epistatic interactions at the level of protein function beyond the physical interactions commonly seen in specific epistasis. We define this as ‘allosteric epistasis’. As allosteric proteins evolve toward new function, such as orthologs in different organisms, their inducer specificity changes to adapt to the new environment^24^. Allosteric proteins may accrue mutations during evolution that would simultaneously affect specificities for old and new inducers and these mutations may also change function by affecting the capability of the protein to produce an allosteric response^25,26^. For instance, a mutant with a stronger affinity for the inducer may not necessarily generate greater output in biological function compared to a mutant with weaker affinity. Thus, the relationship between inducer-binding mutations and biological function governs the fitness landscape of an allosteric protein. Despite the ubiquity of allosteric proteins in every major functional class (transcription, post-translation modification, transport), the role of allosteric epistasis in evolution of novel function in allosteric proteins is not well understood.

Here, we integrate functional, structural, and biophysical analysis to characterize allosteric epistasis in an allosteric transcription factor (aTF). Using computation-guided design, we changed the inducer specificity of TtgR, a promiscuous microbial aTF, to respond to one of its native ligands (resveratrol), but not to another (naringenin) by targeting mutations to positions that directly interact with the ligand^27,28^. By reconstructing all sequence pathways connecting the two states (promiscuous and specific), we found that epistatic interactions of two distinct sets of amino acids separately drive the loss of response to naringenin while increasing response to resveratrol. Thus a few, strong epistatic interactions constrain the number of viable sequence pathways, revealing ridges in sequence space that connect wildtype and redesigned TtgR. Crystal structure of the computationally designed mutant shows that one of the mutations contributes to specificity by reshaping the binding pocket to favor resveratrol over naringenin. We found that allosteric epistasis creates unique biophysical and biological functional landscapes, underscoring the nonlinear relationship between local interactions (ligand affinity) and global effects (allosteric response). Our results highlight the functional and evolutionary complexity of allosteric proteins because^28^ pathways can traverse through multiple adaptive landscapes under evolutionary pressure. Our approach provides a general conceptual and methodological framework to investigate allosteric epistasis in allosteric proteins.

### Computational design of ligand specificity switch

We chose TtgR, a ligand-inducible aTF belonging to the diverse TetR-like protein family, as a target for computational engineering of ligand specificity^28^. TtgR is a 1-component transcriptional system and represents the simplest molecular mechanism for converting biophysical interaction between inducer and protein into a complex biological response like transcription^28^. In the uninduced state, TtgR physically obstructs the RNA polymerase by binding to DNA^28^. When induced, ligand-binding allosterically lowers affinity for DNA, thereby allowing transcription^28,29^. Since TtgR is found in a plant-associated microbe (*Pseudomonas putida*), it is induced by multiple plant molecules including resveratrol and naringenin^27^. Thus, TtgR provides a suitable functional backdrop to investigate the role of epistasis in emergence of novel function (ligand specificity) in an allosteric protein^27,30^. To emulate emergence of novel function, we engineered TtgR to respond to resveratrol and not to naringenin.

We used computational design (Rosetta software suite) to engineer TtgR specificity by generating function-switching mutations that directly interact with the ligand^31^. Less directed approaches may yield a specificity switch, but these can also include distal mutations whose effects on ligand affinity will confound our examination of allosteric epistasis^9,32^. Since our goal was to study how local interactions shape global function, computational design was the appropriate tool as *in silico* mutations are chosen based on interaction energies between protein and ligand^33,34^.

To increase resveratrol specificity, we redesigned the ligand-contacting residues for greater affinity for resveratrol, assuming greater affinity may result in greater specificity. Since Rosetta is a structure-based design tool, the absence of a resveratrol-bound TtgR crystal structure made the design task challenging because the correct position of the ligand in the binding pocket was not known *a priori*. Therefore, we generated a set of diverse starting poses (16) by docking resveratrol conformers in different orientations within the binding pocket (Fig. 1). For each starting pose, we redesigned ligand-contacting residues while permitting constrained rigid-body flexibility of the ligand and torsional flexibility of the protein backbone. We computationally generated approximately 19,000 unique TtgR design variants with an average of 5 mutations per variant (Supplementary Fig. 1). The variants were curated based on Rosetta scoring metrics of stability, repulsion, hydrogen bonds, and protein-ligand affinity to yield a final list of approximately 3,500 sequences for experimental testing. We synthesized oligonucleotides encoding all designed variants as a pool of exact chip-DNA sequences (Twist Bioscience Inc).

**Figure 1:**
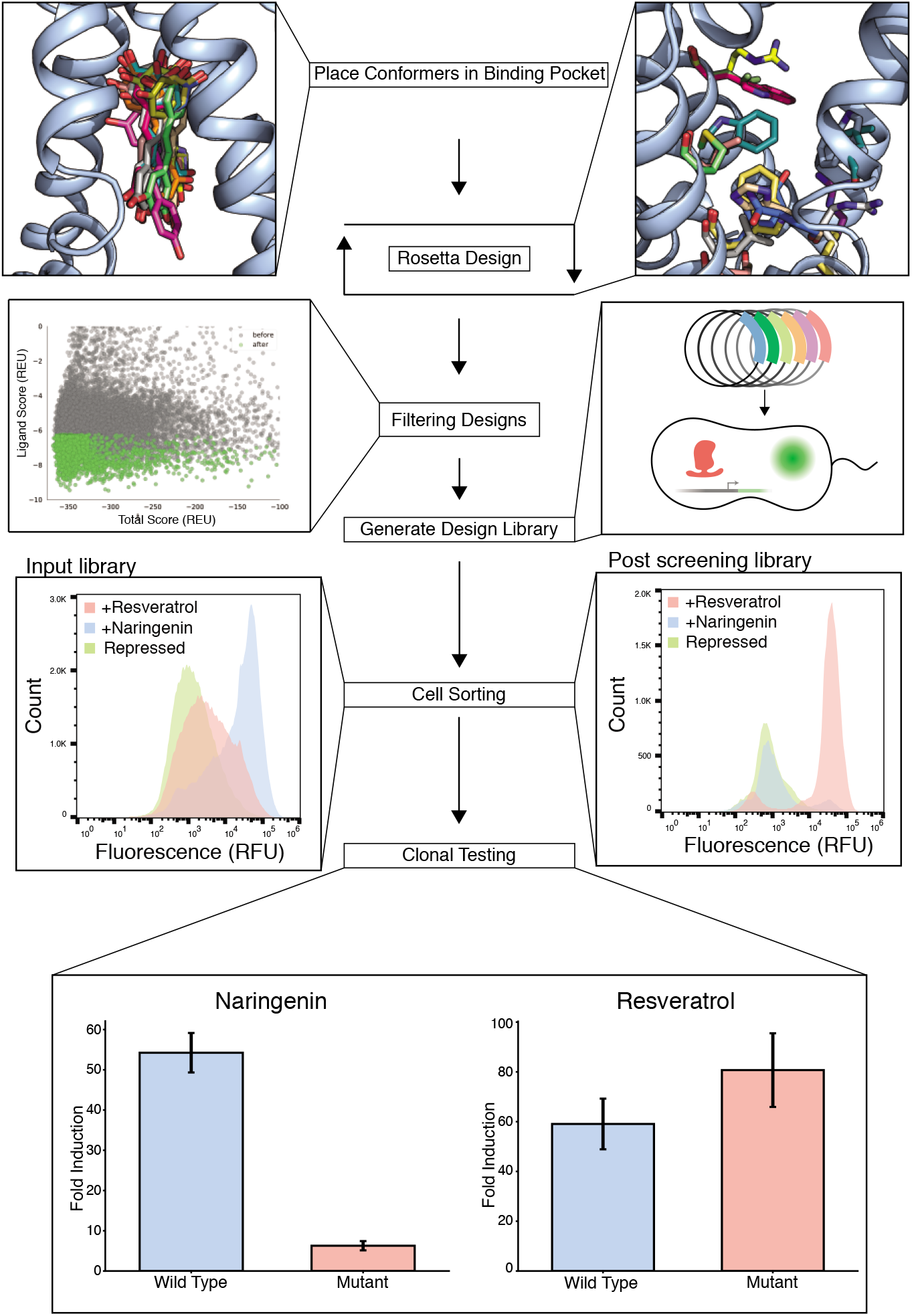
Design of reseveratrol-specific TtgR variant. Resveratrol conformers are docked into TtgR followed by Rosetta-based computational design of the binding pocket. Candidates with favorable Rosetta score metrics (green points) are synthesized and cloned into an expression vector. Distribution of fluorescence in cells containing uninduced TtgR variant library (light green), induced with naringenin (light blue) and resveratrol (red) before sorting (Pre-Sort) and after three rounds of sorting (Post-Sort) are shown. Colony screening identified a quadruple mutant showing resveratrol specificity: C137I/I141W/M167L/F168Y.

To determine the activity of TtgR variants, we designed a pooled screen by sorting *E. coli* cells containing a GFP reporter system regulated by a TtgR operator adapted for *E. coli*. We quantified the activity of variants based on fold induction: the ratio of GFP expression with and without inducer. Fold induction is a simple measure of the transcriptional activity of an aTF that accounts for factors affected by epistasis including DNA affinity, ligand affinity and allostery ^5,8^. The activity of the initial library was greater toward naringenin than resveratrol with a median fold induction of 21-fold and 2.4-fold with naringenin and resveratrol, respectively (Fig 1). To enrich resveratrol-specific variants in the library, we devised a toggled screening scheme where we first sorted variants competent for binding to DNA (low GFP with no resveratrol) followed by sorting variants that can activate expression of the reporter (high GFP with resveratrol) (Supplementary Fig. 2). After three rounds of toggled screening, we observed much greater response to resveratrol than naringenin in the enriched population compared to the input population (Fig.1). From the enriched population, we isolated a resveratrol-specific TtgR variant with four mutations: C137I, I141W, M167L, and F168Y which we will henceforth refer to as the ‘quadruple mutant’. All four mutations were in close proximity to the ligand and no mutations were found elsewhere on TtgR. The quadruple mutant gave 80- and 6-fold induction with resveratrol and naringenin, respectively, compared to 60- and 54-fold of wildtype TtgR (Fig. 1, Supplementary Fig. 3). Therefore, we chose the quadruple mutant as the functional endpoint for characterizing epistasis.

### Epistasis shapes the fitness landscape of resveratrol response

We constructed an experimental fitness landscape to examine epistatic constraints in the transition from wildtype TtgR to the resveratrol specific quadruple mutant. We made all single, double, and triple mutation combinations of the four mutations that provide resveratrol specificity as individual clones, resulting in a total of 16 variants (including endpoints). Experimental fitness landscapes are a useful framework for characterizing epistasis by revealing fitness pathways through mutational intermediates that connect two functional states. Fitness landscapes are commonly illustrated as a series of nodes and edges. Each node is designated by a binary string in which each number corresponds to a mutable position. A zero indicates the wildtype amino acid identity and a one indicates the substituted amino acid. The positions in order from left to right are: 137, 141, 167, and 168 (0000 is wildtype TtgR, 1111 is quadruple mutant, and 0100 represents the I141W mutant). We quantified activity of each variant in terms of fold induction to resveratrol and naringenin. To avoid overestimating epistasis, we applied a log10 transformation to linearize fold induction (Supplementary Fig. 4, Supplementary Fig. 5).

There are 24 possible pathways from wildtype to quadruple mutant (Fig. 2a). We quantified the number of viable pathways in the resveratrol landscape by requiring that each additional mutation must have the same or higher fold induction than its parental variant to emulate neutral drift or progressive gain of function. Viable adaptive pathways from wildtype must go through 0010 as all other single mutants have lower resveratrol response relative to wildtype TtgR (Fig 2a). This restricts the number of available pathways from 24 to a maximum of 6. From 0010, there are three possible double mutants: 0011, 0110 and 1010. Both 0110 and 0011 are not viable as their activity substantially decreases compared to 0010 (Fig 2a). However, 1010 is viable as it gives modestly higher resveratrol response (Fig 2a). Both C137I and M167L manifest as key permissive intermediates in the fitness landscape that allows I141W (1110) or F168Y (1011) to be added. Since 1010 is the only viable double mutant, the number of available pathways between wildtype TtgR and quadruple mutant reduces to two (Fig. 2a, bold red lines). Both triple mutants (1011 and 1110) have higher resveratrol response than 1010 and allow pathways to traverse through sequence space to reach the quadruple mutant, which is the global maxima of this fitness landscape (Fig 2a).

**Figure 2:**
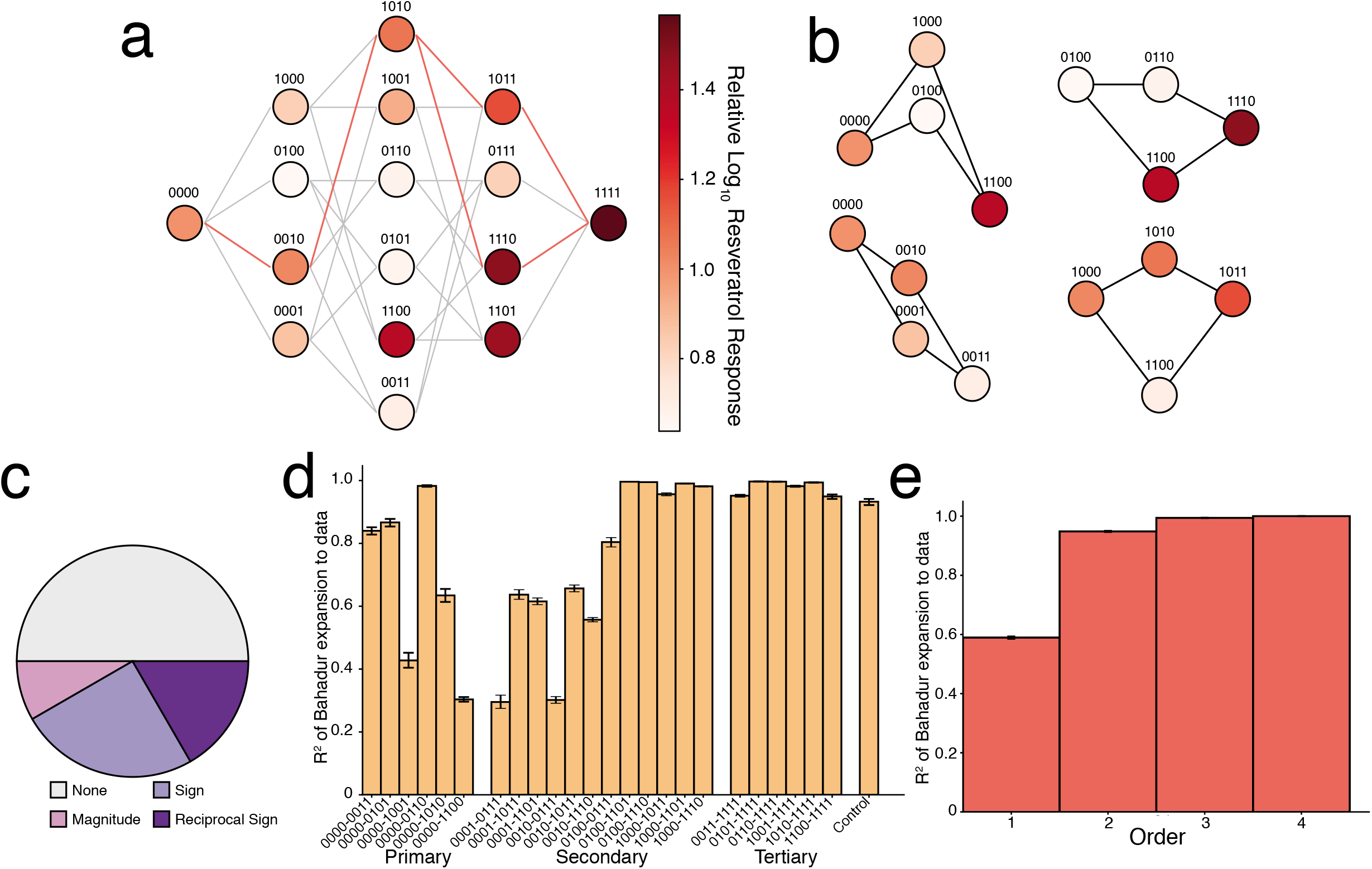
Fitness landscape and epistatic effects for response to resveratrol. **(a)** Fitness landscape of resveratrol response for all 16 TtgR variants connecting wildtype to quadruple mutant with each variant shown as a node in the graph. Each variant is labeled with a binary string corresponding to the presence (1) or absence (0) of a mutation at position 137, 141, 167, or 168 in order. Nodes separated by a single mutation are connected by edges showing viable (bold red) and unviable paths (light gray) through sequence space. Nodes are shaded by log10 fold induction ratio at 250μM resveratrol normalized to log10 fold induction ratio of wildtype TtgR. **(b)** Select subnetworks highlighting key epistatic interactions. **(c)** Different types of epistasis in the 24 subnetworks. **(d)** R^2^ values for each subnetwork individually. Subnetworks are labeled based on the first and last node and are separated into primary, secondary, or tertiary groups. The control group is based off of simulated additive data with standard deviations equivalent to 30% of the mean for each node (Table S1). **(e)** Statistical analysis of the order of epistasis over the entire fitness landscape based on R^2^ values from the Bahadur expansion. The x-axis indicates the highest order terms included in the Bahadur expansion (Order 2 contains 1^st^ and 2^nd^ order terms). Error for (d) and (e) are derived from bootstrap confidence intervals (see methods).

Epistatic interactions are classified as magnitude, sign, or reciprocal sign based on the combined effect of a pair of mutations relative to the effect of each mutant individually. Magnitude epistasis occurs when both mutations individually are beneficial or detrimental and their combined effect is greater in magnitude than sum of their individual effects (Supplementary Fig. 6). Sign epistasis occurs when the effect of one mutation switches from beneficial to deleterious or vice versa depending on if the other mutation is present (Supplementary Fig. 6). Reciprocal sign epistasis occurs when both mutations switch effects when paired (Supplementary Fig. 6).

Two epistatic interactions, C137I-I141W and M167L-F168Y, shape the fitness landscape. The epistatic relationship between I141W and C137I is evident in individual subnetworks (Fig. 2b). A subnetwork contains four variants: a starting background variant, two single mutations added individually, and the combined double mutant. Subnetwork 0000-1000-0100-1100 shows that pairing C137I and I141W creates reciprocal sign epistasis where individually 1000 and 0100 have lower resveratrol response than 0000, but together 1100 has greater response (Fig. 2b). Unless I141W is paired with C137I, it shows drastically lowered resveratrol response (0100, 0110, 0101, and 0111) (Fig. 2Aa. Subnetwork 0100-0110-1100-1110 displays sign epistasis where addition of M167L to I141W results in loss of resveratrol response, but the addition of M167L into 1100 results in an increase in resveratrol response (1110, Fig 2b). The epistatic interaction between M167L and F168Y has differing effects depending on mutational background. In subnetwork 0000-0010-0001-0011, M167L (0010) shows higher resveratrol response and F168Y (0001) shows lowers resveratrol response relative to wildtype (Fig. 2b). Both mutations together (0011) give even lower resveratrol response suggesting prevalence of sign epistasis. In contrast, in the 1000 background M167L-F168Y pair becomes functional (1011) enabling one of the two permissible paths through the 1000-1010-1001-1011 subnetwork (Fig 2b). The increase in resveratrol fold-induction from these two binding-pocket mutations, which are dependent on a preexisting permissive mutation, exemplifies epistasis in allosteric proteins.

While a qualitative description of epistasis is easy to visualize, we wanted to also quantify the extent of epistasis within all individual subnetworks and the entire 16-variant system. We used Bahadur expansion to describe all pairwise and higher order interactions (see methods)^35^. The Bahadur expansion models the activity of the landscape using a linear sum of interaction terms and coefficients. Orders of interactions (first [solo], second [pairwise], third [three way], or fourth [four way]) can be included in this sum to understand their contribution to modeling the behavior of all variants. For each subnetwork, we computed the correlation coefficient between a linear sum of first order interaction terms and actual experimental data. In the simplest case of no epistasis, the correlation coefficient of this comparison (R^2^) is close to 1, but any deviation (R^2^<1.0) indicates prevalence of epistasis. Of the 24 possible subnetworks, 12 are epistatic which includes seven, four, and one instances of sign, reciprocal sign and magnitude epistasis, respectively (Fig 2c). While only half of the subnetworks are epistatic, the location of epistatic subnetworks has a major influence on the shape of the fitness landscape. Therefore, we sought to understand how the strength of epistasis changed with increasing mutations along the fitness landscape. To this end, we classified the subnetworks into primary, secondary and tertiary groups based on whether the originating node has no mutation (0000), a single mutation (1000, 0100, 0010, 0001), or a double mutation (1010, 1001, 0110, 0101, 1100, 0011), respectively (Fig. 2d). Nearly all epistatic interactions exist in the primary and secondary groups including the dominant sign and reciprocal-sign subnetworks (0000-1000-0010-1010 and 0000-1000-0100-1100) (Fig. 2d). Sign and reciprocal sign subnetworks in primary and secondary subnetworks are necessary to create high resveratrol response because most single mutants decrease resveratrol response. Mutations added onto 1010 create non-epistatic or modestly epistatic (magnitude epistasis) interactions that incrementally lead to the fitness peak at the quadruple mutant. Bahadur expansion applied globally (all sixteen mutational combinations) shows that variance between model and experimental data can be explained 60% by first order (linear) terms, 35% by second order (pairwise) terms and third and fourth order epistasis do not significantly contribute to the model (Fig. 2e).

Epistasis thus has a large role in shaping the fitness landscape between the promiscuous wildtype and resveratrol-specific quadruple mutant through key interactions. Although the global expansion first-order terms explain the majority of the variance in the global landscape, higher-order epistatic interactions influence resveratrol response by modulating interactions in secondary and tertiary subnetworks to improve the resveratrol response.

### Epistasis uniquely influences the fitness landscape of each ligand

As inducer specificity changes, the fitness landscape of the same mutational intermediates will differ for each inducer. These differences may reveal alternative adaptive pathways in the fitness landscape of one inducer that circumvent functional ‘dead ends’ in the fitness landscape of another inducer. Therefore, we examined the fitness landscape of naringenin by evaluating naringenin response to all 16 variants to compare with the fitness landscape of resveratrol. We determined the number of viable pathways by requiring that each additional mutation must have same or lower fold induction with naringenin than its parental variant to emulate neutral drift or progressive loss of function (Fig. 3a). None of the 24 possible pathways viably connect wildtype to quadruple mutant because the global minima (variant with lowest naringenin response) in the landscape is the double mutant 0110, not the quadruple mutant (1111) (Fig. 3a). All single mutations decrease naringenin response, but sign epistasis and reciprocal sign epistasis prevent further advance to the quadruple mutant (Fig. 3a). C137I and I141W display strong reciprocal sign epistasis, where I141W alone shows poor naringenin response, but shows strong naringenin response in the background of C137I (see subnetwork 0000-1000-0100-1100, Fig. 3b). A second important interaction in the landscape is between M167L and F168Y (see subnetwork 0000-0010-0001-0011, Fig 3b). These two mutations decrease function in an additive fashion when paired in the wildtype background (0011). However, the same pair of mutations in the 1000 background (1000-1010-1001-1011) creates a series of fitness peaks with high naringenin activity, making paths through these mutations unviable (Fig. 3b). As a result, all steps from 0000 terminate at locations adjacent to detrimental sign and reciprocal sign interactions (Fig 3a).

**Figure 3:**
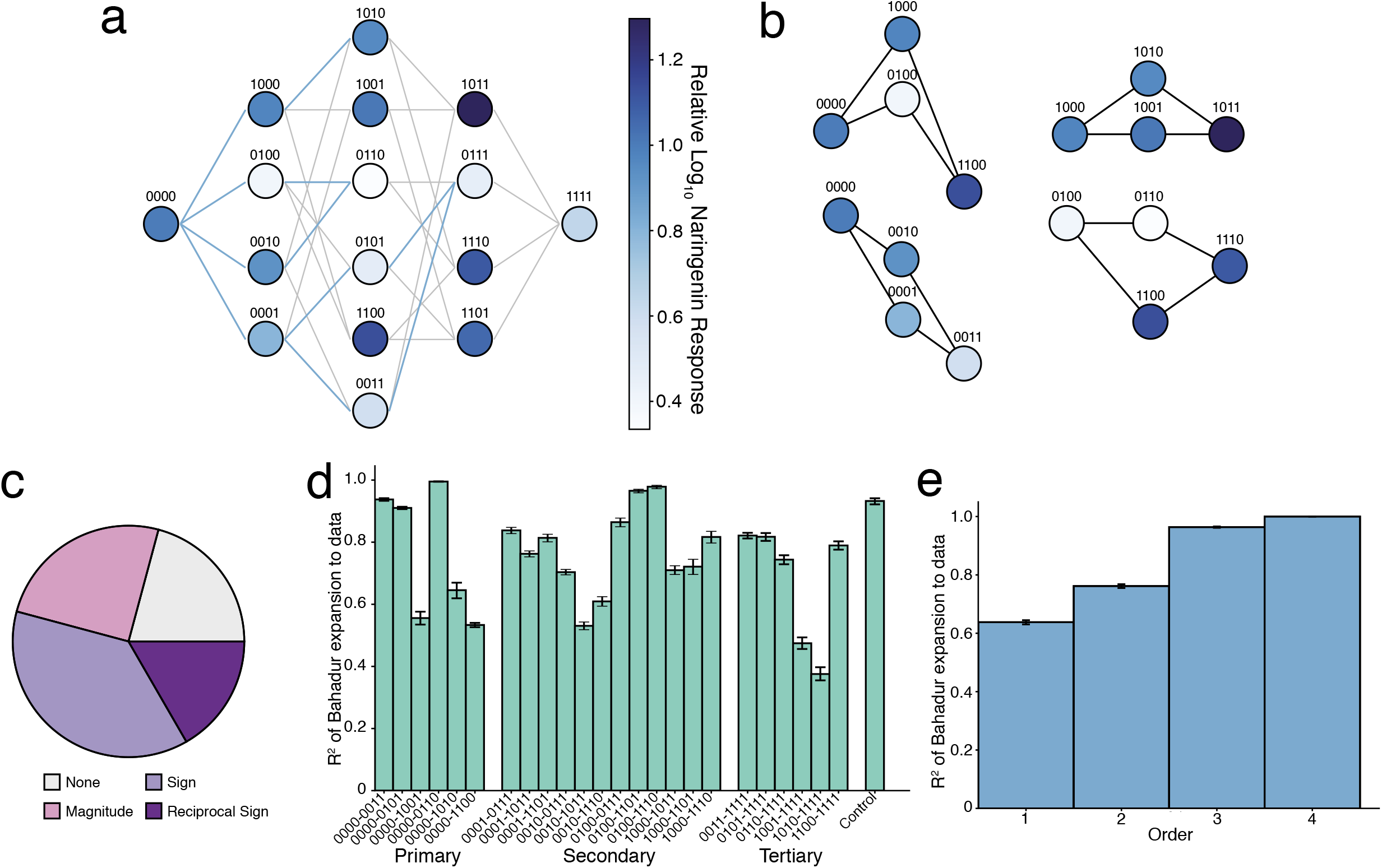
Fitness landscape and epistatic effects for response to naringenin. **(a)** Fitness landscape of naringenin response for all 16 TtgR variants connecting wildtype to quadruple mutant with each variant shown as a node in the graph. Labeling and layout are identical to the resveratrol landscape. Nodes separated by a single mutation are connected by edges showing viable (bold blue) and unviable paths (light gray) through sequence space. Nodes are shaded by log10 fold induction ratio at 2000μM naringenin normalized to log10 fold induction ratio of wildtype TtgR. **(b)** Select subnetworks highlighting key epistatic interactions **(c)** Different types of epistasis in the 24 subnetworks. **(d)** R^2^ values for each subnetwork individually quantified using the Bahadur expansion. Labels, grouping, and the control set are identical to 2d. **(e)** Statistical analysis of the order of epistasis over the entire fitness landscape based on R^2^ values from the Bahadur expansion. Labeling is identical to 2e.

Next, we quantified epistasis both within all individual subnetworks and the entire 16-variant system. Epistasis was much more prevalent in the subnetworks of the fitness landscape of naringenin than those in the fitness landscape of resveratrol. Of the 19 subnetworks (out of 24 total) showing epistasis (Fig. 3c), nine were sign, six magnitude, and four reciprocal sign (Fig. 3c). Three of the six primary subnetworks show no epistasis and each single mutant (1000, 0100, 0010, and 0001) has lower naringenin response compared to wildtype (Fig. 3a,d). However, strong instances of sign and reciprocal sign epistasis in the following subnetworks prevent access to the quadruple mutant: 0000-1000-0010-1010, 0000-1000-0001-1001, and 0000-1000-0100-1100 (Fig. 3d). The secondary subnetworks show predominantly sign and magnitude epistasis, creating ridges of function through the fitness landscape that encompass 1001, 1011, 1100, 1110, and 1101 (Fig. 3e). These regions of high fitness are contrasted by the low naringenin response of the I141W-containing variants, leading to the prevalence of sign epistasis (Fig. 3c, d). The magnitude epistasis subnetworks create subtle changes in naringenin response that propagate the differences created through the sign and reciprocal sign interactions (0001-0011-0101-0111 and 1000-1010-1100-1110) (Fig. 3d). In the tertiary subnetworks, reciprocal sign epistasis exists due to high-functioning triple mutants 1011, 1110, and 1101 rapidly losing function with the addition of the fourth mutation (Fig. 3d). We also quantified the epistasis in the entire system using a Bahadur expansion on all four mutable positions. First-order terms account for 43% of the variation between the model and the data (Fig. 3e). Second-order terms and third-order terms explain an additional 20% and 35% of the variance, respectively. The contribution of fourth order terms is negligible (Fig. 3e).

Epistasis shapes the fitness landscape of each function (naringenin and resveratrol) in distinct ways. The naringenin landscape is roughly the “inverse” of the resveratrol landscape. In the resveratrol landscape, only half of the subnetworks are epistatic and early epistatic networks and low order epistasis create a strong resveratrol response that is propagated through the landscape to the quadruple mutant through additive interactions. However, in the naringenin landscape, the majority of subnetworks are epistatic and higher order epistasis has a large influence on naringenin function through the creation of peaks in the fitness landscape (1011, 1110, and 1101). Strong epistatic interactions are required to create the drastic decrease in response of the quadruple mutant from these peaks.

### Crystal structure reveals molecular basis of specificity of quadruple mutant

To understand the structural basis of TtgR-ligand interactions, we solved high-resolution crystal structures of quadruple mutant (resveratrol-bound and apo) and wildtype TtgR (resveratrol-bound) at a resolution of 1.9Å or better (Table S2). TtgR is a compact, dimeric, all-helical transcription factor with a large cavity between five angled helices forming the ligand binding pocket (Supplementary Fig. 7a,b). The quadruple mutant bound to resveratrol (PDB: 7KD8) is structurally very similar to the wildtype with an all-atom RMSD of 1.2Å over the entire structure. The position and orientation of resveratrol in the wildtype TtgR structure (PDB: 7K1C) resembles the position and orientation of naringenin in a previously solved co-crystal structure of TtgR (PDB: 2UXU)^27^. In both structures, the ligands bind in a vertical mode such that the plane of the molecule is roughly perpendicular to DNA (Supplementary Fig. 7c). In wildtype TtgR, the four mutated positions (C137, I141, M167 and F168) are located approximately in the center of the binding pocket and make nonspecific van der Waals interactions with resveratrol (Fig. 4a, upper panel). Other neighboring residues N110, D172 and H114 make specific hydrogen bonds that stabilize resveratrol in the vertical orientation (Fig. 4a, lower panel). Although both naringenin and resveratrol bind in the vertical orientation, only N110 is able to make a hydrogen bond with both naringenin and resveratrol^27^. The ability of wildtype TtgR to bind multiple ligands likely arises from the nonspecific interactions made by the nonpolar amino acids in the binding pocket.

**Figure 4:**
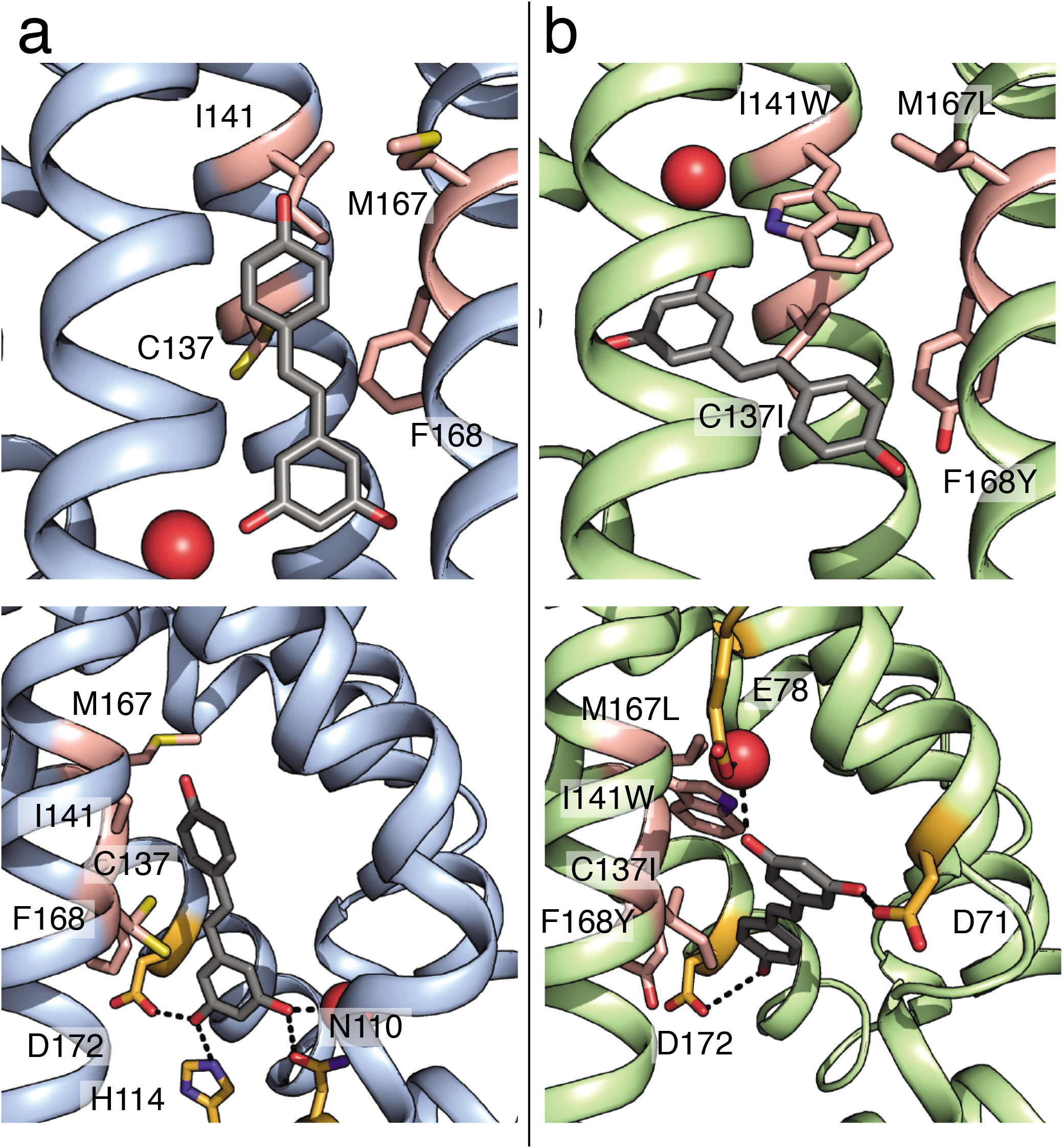
Structural basis for ligand specificity. Wildtype TtgR and quadruple mutant are shown in blue and green ribbons, respectively. Positions 137, 141, 167, and 168 are colored in pink. Resveratrol is shown as gray sticks. **(a)** Binding pocket of resveratrol-bound wildtype TtgR (PDB ID: 7K1C) (upper panel) with residues making hydrogen bonds to resveratrol highlighted in orange (lower panel). **(b)** Binding pocket of resveratrol-bound quadruple mutant TtgR (PDB ID: 7KD8) (upper panel) with residues making hydrogen bonds to resveratrol highlighted in orange (lower panel).

Structure of the quadruple mutant reveals the role of individual residues in ligand specificity. I141W, a mutation critical for resveratrol specificity, creates a large steric barrier that decreases the pocket volume and obstructs the vertical binding orientation of ligands (Fig. 4b, upper panel). Resveratrol is accommodated in the binding pocket in a horizontal binding orientation almost parallel to the plane of the tryptophan. Unlike I141W which plays a clear steric role, the other three mutations (C137I, M167L and F168Y) have a more subtle effect in reshaping the binding pocket through nonpolar interactions. C137I mutation creates a protrusion in the binding pocket that increases shape complementarity to resveratrol (Supplementary Fig. 8a). M167L is buried between the residues in the binding pocket and the dimerization helix and may play a role in positioning the I141W tryptophan to stabilize its horizontal orientation through van der Waals interactions (Supplementary Fig. 8b). F168Y allows the formation of multiple hydrogen bonds with nearby water molecules and may serve to stabilize the structure (Supplementary Fig. 8b). A different hydrogen bonding network consisting of D71, R75, and E78 make hydrogen bonds with the resveratrol molecules in chain A (Supplementary Fig. 8c) and D71, E78, D172, and a nearby water molecule make a hydrogen bond with the single resveratrol molecule in chain B (Fig. 4b, lower panel). Differences in the hydrogen bond network between the wildtype and the quadruple mutant could explain the differences in ligand responsiveness and allosteric activation (fold induction) between them.

Although resveratrol and naringenin share similar chemical backbones, naringenin is bulkier than resveratrol due to the fused carbon rings of the chromanone. This reduces shape complementarity of naringenin to the redesigned binding pocket (Supplementary Fig. 9). The 4-hydroxyphenyl moiety and the carbonyl group of the 4-chromanone backbone of naringenin could create steric clashes with residues lining the wall of the pocket and cause the ligand to sample less space in the pocket compared to resveratrol, which provides a reasonable structural basis for ligand specificity.

The structural basis of ligand specificity relies on the I141W substitution to create a steric barrier to prevent binding in the vertical orientation, which is observed in wildtype TtgR for multiple ligands. In the novel horizontal mode, other ligands may be occluded from the pocket through steric clashes with wildtype residues in the pocket. The epistatic interactions observed in the fitness landscapes for naringenin and resveratrol can be rationalized through examination of the structure. The C137I-I141W pair increases shape complementarity to resveratrol while M167L-F186Y contact the dimerization helix and potentially affect the positioning of nearby residues that interact with the ligand.

### Relationship between biophysical affinity and biological response

Ligand response of an aTF is a complex combination of both biophysical interactions and allostery. Mutations that affect aTF fold induction can do so by altering ligand affinity, DNA affinity, or the allosteric signal upon ligand binding. Since all four mutations are localized to the binding pocket, the observed changes in fold induction of TtgR are likely due to altered binding affinity to ligand, transmission of allosteric signal, or both. To understand the relationship between biophysical affinity and biological response, we compared changes in ligand affinity (Kd) to changes in ligand sensitivity (EC_50_) for both naringenin and resveratrol. We chose mutants in the 0000-1000-0100-1100 subnetwork because it is important for the high resveratrol response in the quadruple mutant. Further, this network shows a strong manifestation of epistasis through reciprocal sign change and is therefore a good model to understand the relationship between biophysical affinity and biological response. We estimated ligand affinity using isothermal titration calorimetry of purified proteins and ligand sensitivity from dose-response curves. Ligand sensitivity is derived from reporter expression and is thus a combination of both allostery and affinity.

Affinity and sensitivity of resveratrol for different variants are generally concordant for resveratrol, with the exception of 1100 (Fig. 5a). As mutations accumulate from wildtype, the affinity and sensitivity generally decrease, suggesting a weaker allosteric response is associated with weaker binding (Fig. 5a). However, although 1100 has similar affinity for resveratrol to 0000 and 1000, its sensitivity is significantly weaker (Fig. 5a). The discordance between affinity and sensitivity is much greater for naringenin than resveratrol. In the case of naringenin, no relationship was evident between affinity and sensitivity across the subnetwork (Fig. 5b). Variant 0100 shows weak affinity, but strong sensitivity for naringenin. In comparison, 1100 shows the opposite: strong affinity, but weak sensitivity for naringenin (Fig. 5b). Although the quadruple mutant has higher resveratrol fold induction than wildtype (Fig.1), it’s affinity and sensitivity for resveratrol is lower than that of wildtype (Fig. 5a). In essence, these examples illustrate the complex relationship between local interactions and their global effects in allosteric proteins.

**Figure 5:**
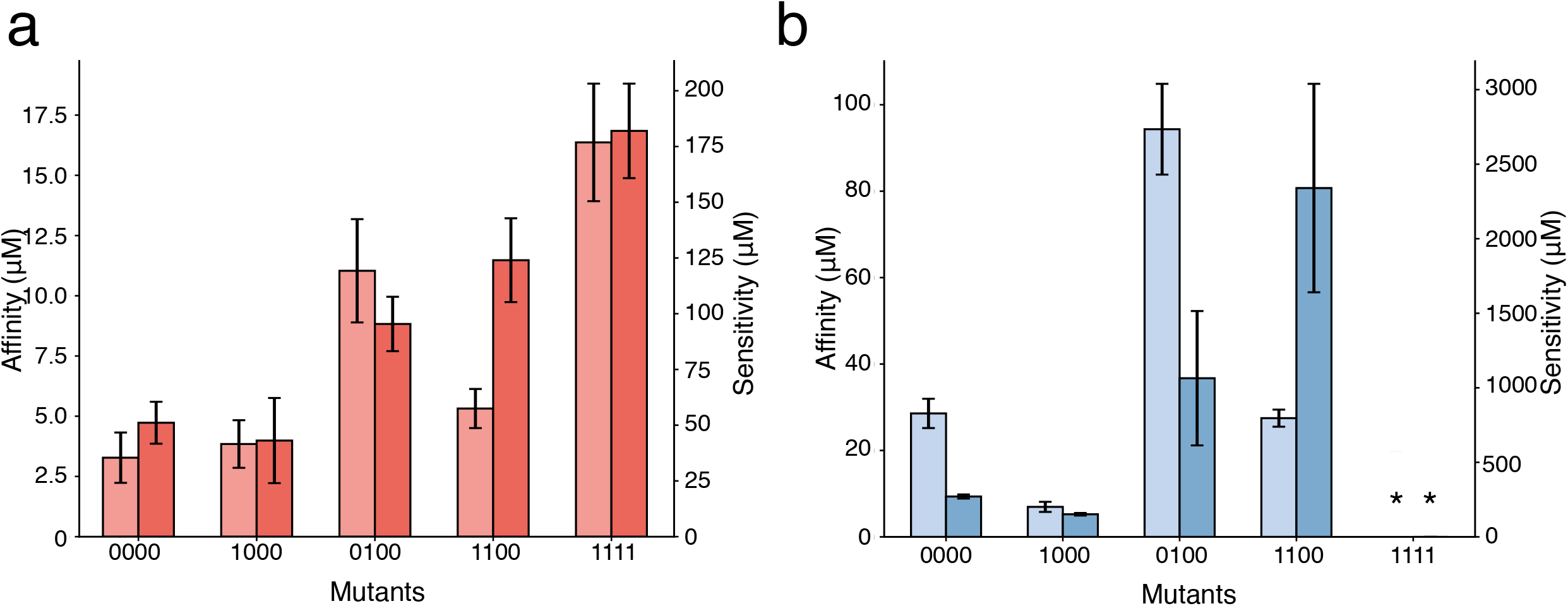
Comparison of biophysical and biological properties of TtgR variants. Ligand affinity (light bar) and EC_50_ sensitivity (dark bar) for resveratrol (red) and naringenin (blue) are shown for TtgR variants 0000, 1000, 0100, 1100, and 1111. Ligand affinity was determined by isothermal calorimetry and EC_50_ sensitivity from fitting dose response curves to the Hill equation. EC_50_ values and error are calculated based on fitting to triplicate dose response curves. ITC values and error are generated from a one-site binding model (see methods).

The 0000-1000-0100-1100 subnetwork displays a unique, ligand-specific pattern of epistasis for biophysical and biological parameters. The mutations we introduced into TtgR have a clear effect on allostery changes in EC_50_ as the complexities of function cannot be simply explained by changes in biophysical affinity. These measurements also suggest that by optimizing a particular protein function (fold induction), other parameters (sensitivity or affinity) may not necessarily stay at fitness maxima as the 1111 mutant shows poor sensitivity to both ligands.

## Discussion

Our study used a constrained set of mutations chosen through *in silico* selection as opposed to natural mutations found in *bona fide* evolutionary pathways. An evolutionary process may have selected a different set of mutations to confer the same functional outcome, leading to the presence of a different pattern of epistasis for either naringenin or resveratrol response. Often in natural evolution, mutations that are distal to the site of interest have a profound effect on protein function^7,20^. These background mutations complicate any examination of key mutations within the targeted area of the protein and their influence on protein function. However, our approach enabled us to examine the propensity of epistasis in a constrained setting where mutations are limited to those that interact directly with the ligand, enabling the examination of the intersection of mutation, biophysical epistasis, and biological epistasis.

In this study, we describe allosteric epistasis, where local interactions between protein and ligand drive global protein function through allosteric communication. By leveraging computational protein design, we engineered four specific local interactions into TtgR, a promiscuous transcription factor that can normally bind to resveratrol and naringenin, to only bind to resveratrol. By characterizing the functional response to both resveratrol and naringenin across all combinations of mutations, we show that the extent of epistasis between mutations affecting global protein function is specific for each ligand. For instance, 50% of subnetworks meet the criteria for epistasis for resveratrol response while 83% of subnetworks are epistatic for naringenin response. However, the fitness landscape of both ligands is shaped by common critical pairs of epistatic interactions (C137I and I141W or M167L and F168Y), though their behavior may be different depending on the context. Biological effects of these mutations are further validated by the crystal structures. The four mutations localize to one face of the binding pocket, making nonpolar interactions with the ligand. C137I and I141W increase shape complementarity of the pocket for resveratrol, but only in an alternative horizontal binding pose. Finally, we highlight that allostery is crucial for modulating biological function beyond residue-level biophysical interactions by examining the differences in sensitivity (EC_50_) and affinity (K_d_).

The multidimensional nature of epistasis in this functional background is a product of the complexity of transcription factor function. TtgR is an allosteric protein; a network of residue interactions facilitates changing DNA affinity in response to ligand binding^30^. By perturbing the amino acid identities in the binding pocket to create a resveratrol-specific mutant, we may also inadvertently affect the allosteric response by disrupting the network of interactions responsible for communicating the ligand binding event to the DNA binding domain. Allosteric epistasis creates additional nonlinear effects on function that, when perturbed, affect both biological and biophysical properties independently.

Our results highlight the dependence of epistasis on protein function (resveratrol or naringenin response) and the prevalence of distinctive adaptive landscapes between functions within the same set of mutations. These landscapes can become interconnected by changing selection pressures between different protein functions. On an evolutionary scale, simultaneously changing protein sequence and selection pressure may enable improbable trajectories by bypassing epistatic barriers to reach previously inaccessible mutational states. In our case, higher order epistasis which prevents access to the quadruple mutant in the naringenin landscape, could be bypassed by toggling between naringenin and resveratrol selection pressures. The evolution of allosteric proteins is inherently dependent on allosteric epistasis and the interactions arising between mutations in these proteins uniquely affects multiple adaptive landscapes.

## Acknowledgements

We thank Dr. Tina Wang, Dr. Phil Romero and Phil Huss for critical review of the manuscript. This work was supported by is partially supported by the US Army Research Office (W911NF-17-1-0043). This research was performed using the compute resources and assistance of the UW-Madison Center for High Throughput Computing (CHTC) in the Department of Computer Sciences. The CHTC is supported by UW-Madison, the Advanced Computing Initiative, the Wisconsin Alumni Research Foundation, the Wisconsin Institutes for Discovery, and the National Science Foundation, and is an active member of the Open Science Grid, which is supported by the National Science Foundation and the U.S. Department of Energy’s Office of Science. This research used resources of the Advanced Photon Source, a U.S. Department of Energy (DOE) Office of Science User Facility operated for the DOE Office of Science by Argonne National Laboratory under Contract No. DE-AC02-06CH11357. Use of the LS-CAT Sector 21 was supported by the Michigan Economic Development Corporation and the Michigan Technology Tri-Corridor (Grant 085P1000817). We thank Spencer Anderson, Joseph Brunzelle, Elena Kondrashkina, and Zdzislaw Wawrzak (LS-CAT) and Craig Ogata, Michael Becker and Nagarajan Venugopalan (GM/CA@APS) for synchrotron beamline support. K.N is supported by NIH National Research Service Award T32 GM07215 and the Robert and Katherine Burris Biochemistry Fund.

## Author contributions

K.N and S.R designed the study, analyzed the data, and wrote the manuscript. K.N performed all experiments. N.H carried out computational design. R.S and C.B purified and crystallized the proteins.

## Methods

### Computational Design

Protein modeling and design was performed with Rosetta version 3.5 (2015.19.57819)^34,36^. Python and shell scripts for generating input from Rosetta and analyzing from Rosetta are available at: https://github.com/raman-lab/biosensor_design

#### Structure and ligand preparation

The high-resolution TtgR structure co-crystalized with tetracycline was selected as the starting point for computational design (PDB ID: 2UXH)^27^. The structure was prepared for use in Rosetta by performing an all-atom, coordinate-constrained relaxation^37^.

#### Commands

Rosetta/main/source/bin/idealize_jd2.linuxgccrelease -database

Rosetta/main/database/ -in::file::fullatom -s 2UXH.pdb -extra_res_fa LG.params - no_optH false -flip_HNQ

Rosetta/main/source/bin/relax.linuxgccrelease -database Rosetta/main/database/ - relax::sequence_file always_constrained_relax_script - constrain_relax_to_native_coords -relax::coord_cst_width 0.25 -relax::coord_cst_stdev 0.25 -s 2UXH_idealized.pdb -in::file::native 2UXH_idealized.pdb -extra_res_fa LG.params -in::file::fullatom -no_optH false -flip_HNQ

Rosetta/main/source/scripts/python/public/molfile_to_params.py -n resveratrol.params - p resveratrol.pdb

#### Protein design simulations

The RosettaScripts protocol used to design the ligand binding pocket of each starting TtgR-resveratrol complex was based on enzyme design protocols^31,38^.

#### Command

Rosetta/main/source/bin/rosetta_scripts.linuxgccrelease -database Rosetta/main/database/ -parser::protocol enzdes.xml -in::file::s 2UXH_resvertrol.pdb - extra_res_fa resv.params -use_input_sc -packing:linmem_ig 10 -ex1-ex2 - run:preserve_header -enzdes_out -enzdes:bb_min_allowed_dev 0.2 - enzdes:loop_bb_min_allowed_dev 0.5 -enzdes:minimize_ligand_torsions 15 - parser::script_vars ligchain=X resfile=TtgR.resfile -out::pdb -nstruct 10

The TtgR.resfile is a plain text file containing the amino acid position numbers that were able to be mutated during design, and these were positions 137, 141, 167, 168, 171, 172, 175, and 176. We used UW-Madison’s Center for High Throughput Computing computer cluster to perform 320,000 different design simulations. The resulting designed structures were curated to yield the set of sequences that we synthesized to isolate resveratrol-specific TtgR variants.

#### Selection of designs for synthesis

We selected computational designs for synthesis by first removing designs that were repetitive and then removing designs that were energetically unfavorable. The criteria for unfavorable energies were selected empirically based on the distribution of energies for all designs to yield approximately 10^4^ sequences for synthesis. Specifically, on each unique design, ΔΔG stability calculations were performed on designed residues to ensure the number of destabilizing changes was limited. If the mutation destabilized the TtgR-resveratrol complex by 0.5 Rosetta Energy Units (REU), the residue was reverted to its wild-type identity. After this, non-unique designs were again removed. The unique designs were filtered using distance from the median absolute deviation of several salient Rosetta scoring metrics including total ligand binding energy, hydrogen bond energy, Leonard-Jones repulsive energy, solvation energy, and total score, which is a weighted, linear combination of all score terms in the energy function^33^. Designs that passed this filter were synthesized for library screening.

#### Commands

./biosensor_design/fas_from_pdb_stdout.py *.pdb > TtgR_resveratrol_all_designs.fasta

./biosensor_design/uniquify_fas.py TtgR_resveratrol_all_designs.fasta > TtgR_resveratrol_unique_designs.fasta

./ddg_monomer.static.linuxgccrelease -database./database @ddg_flags -in:file:s design_pdb.pdb -ddg::mut_file list_of_positions_to_calc_ddg.mutfile -ddg::iterations 50

./gen_enzdes_cutoffs.py concatentated_design_score_file.sc -c median_abolute_deviation_cutoffs.txt -o designs_passing_filter.sc

#### The median absolute deviation cutoffs used were

total_score < +1 MAD
fa_rep < +3 MAD
hbond_sc < +3 MAD
tot_burunsat_pm < +3 MAD
%(LIG)s_fa_rep < +3 SD
%(LIG)s_hbond_sc < +3 MAD
%(LIG)s_burunsat_pm < 2.5 ABS
%(LIG)s_total_score < −1 MAD

### Library synthesis

#### Creating sfGFP reporter plasmid

The sfGFP reporter plasmid was constructed using a backbone containing the ColE1 origin and a kanamycin resistance gene. The TtgR operator sequence was modified to contain canonical −10 (5’-TATAAT-3’) and −35 (5’-TTGACA-3’) elements in the promoter. A strong RBS (g10) was chosen for high sfGFP expression^39^. The TtgR operator-RBS sequence was constructed via sequential PCR reactions with overlapping primers containing homology to the pColE1 backbone 5’ of sfGFP. The plasmid was annealed using isothermal assembly using 0.16pmol of backbone and 0.43pmol of promoter^40^. DH10B cells (NEB) were transformed with the pColE1 reporter plasmid and plated on LB-kanamycin agar (50μg/mL). A colony was selected and grown in LB-kanamycin media (50μg/mL) shaking for 16 hours at 37°C. An aliquot of the culture was stored at −80°C in 25% glycerol. Plasmids were isolated using a DNA miniprep kit (Omega BioTek) according to the manufacturer’s protocol. The insertion of TtgR operator sequence was confirmed via Sanger sequencing.

#### Creating TtgR expression plasmid

The TtgR expression plasmid used the SC101 origin and a spectinomycin resistance gene. The constitutive promoter-RBS combination apFAB61-BBa_J61132 and the TtgR gene were amplified via KAPA HiFi PCR mix (Roche) using primers with homology to the pSC101 backbone^41^. The TtgR-pSC101 construct was generated using isothermal assembly (0.046pmol backbone and 0.24pmol TtgR) and DH10B cells were transformed with the TtgR-pSC101 construct. A colony was selected and grown in LB-spectinomycin media (50μg/mL) shaking for 16 hours at 37°C. An aliquot was stored at −80°C and plasmids were isolated and verified as described previously.

#### Library cloning

Rosetta-designed sequences were synthesized as exact oligos (Twist Biosciences). Oligos were converted to double-strand DNA using qPCR and purified on a spin column (EZNA Cycle Pure kit from Omega BioTek). The pSC101 backbone was amplified with two separate primer pairs encoding BsaI cut sites that matched the insertion location of the oligos on the TtgR gene. The amplified backbone was treated with Dpn1 for 16 hours at 37°C (NEB) followed by a purification using a spin column. The backbone was treated with BsaI (NEB) for 2.5 hours at 37°C followed by purification using a spin column. The digested backbone was treated with Antarctic phosphatase (NEB) for 1 hour at 37°C followed by purification using a spin column. A golden gate reaction (NEB) was performed using 0.12pmol backbone and 0.89pmol library oligo in roughly a 1:7 molar ratio and incubating for 30 cycles of 37°C for 5min and 16°C for 5 min followed by 60°C for 5min. A control reaction was made using just the pSC101 backbone with no Rosetta oligos added. The golden gate reactions were dialyzed using semi-permeable membranes (Millipore) for 1 hour at 25°C against dH2O. 25μL of C3020 cells (NEB) were transformed with 2μL of the dialyzed golden gate mixture via electroporation. Cells recovered for 1 hour in SOC media shaking at 37°C and were diluted 5X with LB. Dilutions of 100X, 500X, and 1,000X were plated to calculate transformation efficiency relative to the control. A transformation was considered successful when CFU/mL of the Golden Gate reactions exceeded CFU/mL of control reactions by a factor of 10 or more. Cells grew for 6 hours post transformation before the culture was diluted 50X and grown overnight shaking at 37°C for 16 hours. Plasmids of the library were harvested using a DNA miniprep kit and stored at −20°C.

#### Preparing electrocompetent cells with reporter plasmid

An aliquot of the pColE1 frozen stock was streaked on a LB-kanamycin agar plate and grown for 16 hours at 37°C. A single colony was selected and grown in LB-kanamycin media shaking for 16 hours at 37°C. The culture was diluted 50X and grown at 37°C to an OD_600_ of 0.6. Cells were placed on ice and 5mL aliquots were centrifuged at *5,500g* for 5 minutes at 4°C. Pellets were resuspended, washed with ice cold dH2O, and spun at *5,500g* twice. The cells were resuspended in 20μL of water to create electrocompetent DH10B containing the pColE1 plasmid. DH10B *E.coli* containing the pColE1 reporter plasmid were transformed with the initial Rosetta library in pSC101 via electroporation. The transformed cells were recovered for 1 hour shaking at 37°C before dilutions were plated on LB-kanamycin/spectinomycin agar plates (50μg/mL each) to calculate transformation efficiency. The remaining cells were diluted 5X with LB-kanamycin/spectinomycin media and grown shaking at 37°C for 16 hours. A frozen stock was made with 25% glycerol.

#### Sorting the resveratrol library

50μL aliquots of the co-transformed Rosetta libraries were thawed on ice and inoculated into 5mL of LB-kanamycin/spectinomycin and grown shaking at 37°C to an OD_600_ of 0.2. Wildtype co-transformed TtgR sensor+reporter was also inoculated as a reference. These were then split into 4 1mL aliquots and inoculated with either 500μM naringenin (DMSO), 95μM resveratrol (ethanol), DMSO, ethanol and grown for 14 hours at 37°C shaking. Cells were diluted 50X in ice cold PBS (137mM NaCl, 2.7mM KCl, 10mM Na_2_HPO_4_, 1.8mM KH_2_PO_4_) and stored on ice prior to sorting.

Sorting was conducted using a Sony SH800 cell sorter. Cells were excited by a 488nm laser and GFP fluorescence was captured through a 525/50 filter. Gain settings were adjusted such that all cells fell between 10^2^ and 10^6^ RFU. 100,000 event measurements of all libraries, induced and repressed, were taken to draw gates according to population percentage.

Sorting followed an induced-repressed schema; the first library sort consists of taking 500,000 cells of median 50% of fluorescence from the nontreated distribution. This sort isolates cells that contain TtgR variants capable of repressing GFP expression. Cells were sorted into 2mL of LB. LB as added to a final volume of 5mL and incubated for 1 hour at 37°C shaking. Kanamycin and spectinomycin were added after 1 hour to a final concentration of 50μg/mL each from 1mg/mL stocks. These grew to an OD_600_ of 0.2 before frozen stocks were made in 25% glycerol. A small aliquot was stored as a frozen stock at −80°C in 25% glycerol. The remaining culture was induced with naringenin, resveratrol, DMSO, or ethanol at an OD_600_ of 0.2.

The next sort consisted of isolating 100,000 cells in the top 5% of fluorescence from the resveratrol-induced library. This subpopulation was grown as described previously and induced with 95μM resveratrol at an OD_600_ of 0.2. The final sort consisted of isolating 500,000 cells the bottom 60% of the nontreated fluorescence distribution. The sorted cells were incubated at 37°C until the culture reached an OD_600_ of 0.2. A frozen stock was stored at −80°C in 25% glycerol.

#### Clonal Testing

Aliquots of the sorted library, wildtype TtgR, and a GFP-positive control were thawed on ice. 50μL of the library was plated on LB-kanamycin/spectinomycin and incubated at 37°C for 16 hours. The GFP control aliquot was streaked on LB-kanamycin and the wildtype TtgR aliquot was streaked on LB-kanamycin/spectinomycin and incubated in the same fashion. Colonies were selected from each plate and inoculated into 150μL of LB in a 96 well plate. The colonies were incubated at 37C shaking in a SBT1500-H microplate shaker (Southwest Science) and grew to saturation (approximately 8 hours). The cultures were diluted 15X into fresh LB with either 500μM naringenin or 95μM resveratrol and incubated in a Synergy HTX plate reader (BioTek) for 16 hours at 37°C. The performance of each colony was measured using the ratio of fluorescence to optical density (RFU/OD_600_). The ratio of this measurement in the presence and absence of ligand defined the response to each ligand. Successful colonies had higher response for resveratrol than for naringenin. These colonies were sequenced using Sanger sequencing.

### Testing of combinatorial mutants

#### Generation of combinatorial mutants

The 14 mutational intermediates were generated using eight primers specifically encoding combinations of either 137+141 or 167+168. The resulting oligos were inserted into the TtgR-pSC101 plasmid using isothermal assembly using .042pmol of backbone and 0.8pmol TtgR. DH10B *E.coli* cells (NEB) were transformed with the resulting reaction via electroporation. Colonies were selected and sequenced to verify the correct mutations were present. The correct colonies were inoculated into LB-spectinomycin and incubated at 37°C for 16 hours. An aliquot was stored at −80°C in 25% glycerol and plasmids were harvested from the remaining culture. DH10B cells were cotransformed with the 14 TtgR-pSC101 plasmids and the pColE1 reporter plasmid. These were grown for 16 hours shaking at 37°C in LB-kanamycin/spectinomycin media and frozen in 25% glycerol at - 80C.

#### Dose response curves

A 250mM stock of naringenin was made in DMSO and a 100mM stock of resveratrol was made in ethanol. The TtgR-pSC101/pColE1 frozen stocks were struck out onto LB-kanamycin/spectinomycin plates. Colonies were selected and inoculated into 150uL LB in a 96-well plate. These grew in a microplate shaker to saturation (approximately 8 hours) at 37°C. The cultures were diluted 15X into fresh LB-kanamycin/spectinomycin in a 96-well plate with varying concentrations of either naringenin (0μM, 10μM, 25μM, 50μM, 75μM, 100μM, 250μM, 500μM, 750μM, 1000μM, 1500μM, 2000μM) or resveratrol (0μM, 2.5μM, 5μM, 7.5μM, 10μM, 25μM, 50μM, 75μM, 100μM, 150μM, 200μM, 250μM). A series of naringenin and resveratrol stock concentrations were made such that a 50X or a 100X dilution, respectively, would yield the desired concentrations in the assay. Most variants were assayed with three biological replicates. Variants with more biological noise (1010, 1001, 1110, and 1101 for naringenin and 1001, 1000, 0001, and 0011 for resveratrol) were assayed with six replicates. The assay was incubated in the microplate shaker for 14 hours at 37°C shaking. Cells containing wildtype TtgR pSC101 with the pColE1 reporter and cells containing pColE1 reporter alone served as controls and were included on every plate. A set of 6 biological replicates of a sfGFP positive control were induced with both sets of ligands and concentrations.

Cells were diluted 50X in ice cold PBS. Fluorescence measurements were conducted on a LSR-Fortessa system (BD Biosciences) using a 488nm laser for excitation and a 530/30 filter for fluorescence emission. Using gates on FSC-H vs FSC-A, 100,000 events were gathered per well. To account for changes in fluorescence that are independent of TtgR function, raw fluorescence values were normalized by fold changes in sfGFP fluorescence in the positive control (N=6). The median values of the fluorescence distributions were used as the basis for fold induction calculations. Fold induction as calculated by obtaining the ratio of induced average median fluorescence to baseline average median fluorescence.

### Quantifying epistasis

#### Analyzing fluorescence data

The mean and standard deviation of each concentration of ligand for each combinatorial mutant were used to calculate a fit using the Hill equation as a function of ligand concentration (x)^42^.

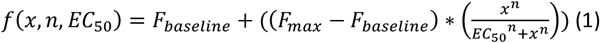

TtgR function was defined as the maximum fold induction of the system, which is the ratio of the median fluorescence at the highest ligand concentration and the median fluorescence at 0μM ligand:

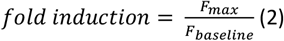

The Python 2.7 function curve_fit() from the Scipy module was used to fit the dose response curves to the Hill equation (Supplementary Fig. 10, Supplementary Fig. 11)^43^. This function provides both fit parameters and error as a covariance matrix as output.

#### Bahadur expansion

The Bahadur expansion was used to analyze the data^35^. Each mutant can be represented as a numerical string (z string), where each mutable position is one number (zi) in the string. A wild type residue at a position is designated by a −1 while the mutated residue is designated by a 1. The mutant M167L+F168Y thus becomes [−1, −1, 1, 1]. The interaction terms can be modeled as follows:

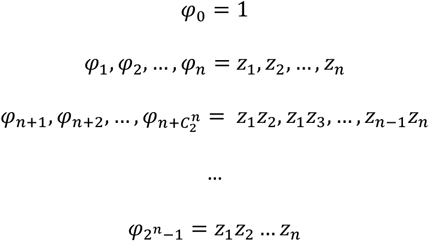

An orthonormal matrix of psi-values is created based on the combinations of mutations within the set (Supplementary Table 3). The Bahadur coefficients can be calculated using this orthonormal matrix and a fluorescence values f(x) for a particular mutant x in the set of all mutants X.

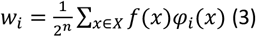

The fluorescence of each combinatorial mutant can be calculated based on the Bahadur coefficients and z string.

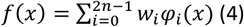

The R^2^ between the modeled fluorescence values and the experimental data is 1.0 when all interaction terms are included in the expansion. By truncating Eq. (4) to contain only low-order interactions, the effect of these contributions to the model can be determined. The expansion was applied to the full set of mutations (4 positions) and modeled using first order terms; first and second order terms; first, second, and third order terms; and all terms (Supplementary Fig. 12). An identical approach was applied to all 24 subnetworks and utilized only first order terms in the reconstruction (Supplementary Fig 13).

Errors in the R^2^ statistics were estimated using a Monte Carlo simulation. 500 sets of fluorescence values for all mutants were sampled based on experimental fluorescence means and standard deviations following a Gaussian distribution using the NumPy module in Python 2.7^44,45^. Eq. (3) and (4) were applied to reconstruct the fluorescence values and calculate R^2^ values between the sampled model and the sampled data to give a distribution of R^2^ values. Bias-corrected adjusted confidence intervals were calculated by obtaining the average R^2^ of 10,000 bootstrap iterations of the Monte Carlo simulation R^2^.

A control set of additive data was used to calculate the R^2^ of data showing no epistasis (Supplementary Table 1). This subnetwork was analyzed using the same approach as the subnetwork workflow.

### Protein characterization

#### Purifying proteins for isothermal titration calorimetry

The TtgR gene for variants 0000, 1000, 0100, 1100, and 1111 were cloned into a pET31B vector downstream of the T7 promoter for lac-inducible transcription control using isothermal assembly with 0.18pmol backbone and 0.392pmol TtgR. MBP was amplified with primers to add a C-terminal His-tag and TEV site and inserted into the TtgR-pET31 B vector upstream of TtgR to create a MBP-His-TtgR fusion with a TEV cleavage site between the His-tag and the TtgR protein. BL21 chemically competent cells (NEB) were transformed with 20ng of pET31B vector. Dilutions of transformants were plated on LB-ampicillin agar. A colony was selected and grown in 5mL LB-ampicillin media shaking at 37°C for 16 hours. This culture was added to 500mL autoinduction media (Terrific Broth, 0.8% glycerol, 2mM MgSO4, 0.375% (w/v) aspartic acid, 0.015% (w/v) glucose, 0.5% (w/v) lactose) and grown for 8 hours at 37°C shaking. The culture was grown for an additional 16 hours at 25°C shaking.

The cells were spun down at 5,500*g* for 15 minutes at 4°C. The supernatant was removed and the cells were resuspended in a lysis buffer (300mM NaCl, 50mM HEPES, 1mM PMSF, 1mg/mL Lysozyme, 5mM BME, 10% glycerol, pH 7.5). A Q500 sonicator (Qsonica) was used to lyse cells using a 5 second on, 15 second off sonication protocol for 4 minutes total sonication time. The lysate was centrifuged at 14,000*g* for 45 minutes at 4°C. The supernatant was isolated and filtered through a 0.22μm filter. The filtered supernatant was purified on an Akta Start using 2 5mL HisTrap HP columns. The column was washed with 5 column volumes (CV) IMAC-A (500mM NaCl, 20mM Imidazole, 20mM MOPS, 0.3mM TCEP, pH 7). MBP-6His-TtgR was eluted with a gradient of 100% IMAC-A to 100% IMAC-B (500mM NaCl, 500mM Imidazole, 20mM MOPS, 0.3mM TCEP, pH7) over 5CV and collected in 2mL fractions. Fractions with the highest absorbance at 280nm (A280) were combined and dialyzed in 8L of dialysis buffer A (100mM NaCl, 20mM MOPS, 0.3mM TCEP, pH 7.5). TEV was added to the proteins prior to dialysis at a ratio of 1:50 w/w TEV:TtgR. Dialysis occurred over a 16 hour interval at 4°C while stirring at low speed. Dialyzed protein was centrifuged at 14,000*g* for 10 minutes at 4°C. The supernatant was passed through a 0.22μm filter and loaded onto the HisTrap columns at 5mL/min. The column was washed with 5CV of IMAC-A and 2mL fractions were collected. 5CV of IMAC-B was used to remove the MBP-6His from the column. The column was washed with an additional 10CV IMAC-A. Wash fractions with high A280 were combined and reapplied to the column. The column was washed with 5CV of IMAC-A and 2mL fractions were collected. 5CV of IMAC-B was used to strip the MBP-6His from the column. Fractions with high A280 were combined and dialyzed in 4L of dialysis buffer C (100mM NaCl, 20mM MOPS, 10mM MgCl2, 0.3mM TCEP, pH 7.8). The protein was centrifuged at 14,000*g* for 10 minutes at 4°C. The supernatant was passed through a 0.22μm filter. The protein was concentrated to approximately 9mg/mL and frozen in 60μL aliquots in liquid nitrogen before storing at −80°C. Dialysis buffer C was passed through a 0.22μm filter and stored at 4°C for ITC experiments.

#### Determining binding affinity of TtgR variants

Stocks of 250mM naringenin and 100mM resveratrol were diluted to 500μM and 250μM, respectively, in dialysis buffer C. Aliquots of TtgR were thawed on ice and diluted to a final concentration of 7.5μM. DMSO or ethanol was added to the TtgR solution to match the solution composition of the naringenin or resveratrol dilutions. An aliquot of dialysis buffer C was also prepared with DMSO or ethanol for a control injection and to wash the sample cell between ITC injections.

The ITC experiments were conducted on a VP-ITC (MicroCal). An initial control injection scheme consisted of loading the sample cell with dialysis buffer C and performing a series of 10 10μL ligand injections with 10 minute intervals at 25°C. The sample cell was washed 5 times with dialysis buffer C before the 7.5μM protein solution was loaded. 25 10μL naringenin injections or 28 10μL resveratrol injections occurred in 10 minute intervals at 25°C.

Data analysis was primarily conducted using Origin 7.0 (MicroCal). The heats of injection from the control sample were averaged. The protein-ligand injection profile was subtracted by this average heat prior to curve fitting. Due to low affinity for both naringenin and resveratrol, the stoichiometry of binding was fixed to 1 to reduce the degrees of freedom prior to fitting. The curves were fit with the single binding site model (Supplementary Fig. 14).

### X-ray crystallography

#### Purifying Proteins for X-ray crystallography

TtgR-pET31B vector was electroporated into BL21 cells (NEB) and recovered in 1mL SOC. The cells were incubated for 1 hour at 37°C before serial dilutions were plated on LB-ampicillin (100μg/mL) plates. A single colony was selected and incubated in 5mL LB-ampicillin (100μg/mL) at 37°C shaking for 3 hours. The 5mL culture was added to 500mL LB-ampicillin media and incubated at 37°C shaking at 250rpm for approximately 3 hours until the OD_600_ reached 0.6. The culture was induced with 100μM IPTG followed by an incubation at 16°C for 16 hours shaking at 250rpm.

The cells were spun down at *5,500g* for 15 minutes at 4C. The supernatant was removed and the cells were resuspended in a lysis buffer (300mM NaCl, 50mM HEPES, 1mM PMSF, 1mg/mL Lysozyme, 5mM BME, 10% glycerol, pH 7.5). A Q500 sonicator (Qsonica) was used to lyse cells using a 25 second on, 50 second off sonication protocol for 3 minutes and 45 seconds total sonication time. The lysate was centrifuged at 14,000*g* for 45 minutes at 4°C. The supernatant was isolated and filtered through a 0.22μm filter. The filtered supernatant was purified on an Akta Start (Cytiva) using a 5mL HisTrap HP columns (Cytiva). The supernatant was loaded onto the column at a flow rate of 5mL/min. The column was washed with 5 column volumes (CV) IMAC-A. MBP-6His-TtgR was eluted with a gradient of 100% IMAC-A to 100% IMAC-B over 10CV and collected in 2mL fractions. Fractions with the highest absorbance at 280nm (A280) were combined and dialyzed in 8L of dialysis buffer A. TEV was added to the proteins prior to dialysis at a ratio of 1:50 w/w TEV:TtgR. Dialysis occurred over a 16 hour interval at 4°C while stirring at low speed.

TtgR was isolated from MBP-6His through a subtractive IMAC protocol using the Akta Start and 5mL HisTrap HP column. The dialyzed protein was centrifuged at 4,000*g* for 10 minutes at 4C. Supernatant was passed through a 0.22μm filter and applied to the HisTrap column at 5mL/min. 5CV IMAC-A was used to wash the column while 2mL fractions were collected. 2.5CV IMAC-B was used to remove the MBP from the column and 5mL fractions were collected. Wash fractions with high A280 were combined and dialyzed in 4L of dialysis buffer B (50mM NaCl, 5mM MOPS, 0.3mM TCEP, pH 7.5). EDTA was added to the protein wash fractions to a final concentration of 10mM prior to dialysis. Dialysis occurred over a 16 hour interval at 4C while stirring at low speed. TtgR was concentrated to 10mg/mL using spin concentrators. Samples were spun at intervals of 3,500*g* for 5 minutes and mixed via pipette between spins. Concentrated TtgR was separated into 60μL aliquots and frozen in liquid nitrogen prior to storage at −80°C.

#### Size exclusion chromatography

Samples of TtgR wildtype and mutant proteins were received frozen in 5 mM MOPS, pH 7.4, 50 mM NaCl, 0.3 mM TCEP. Samples were thawed and centrifuged for 5 minutes at 21,130*g*. Sample supernatants were filtered with a 0.22 micron MillexGV syringe filter unit (Millipore) before applying to an equilibrated 10 mm x 300 mm Superdex 200 column (GE Healthcare). Chromatography was performed on a GE AKTA FPLC system. Column buffer was 20 mM HEPES, pH 7.5, 350 mM NaCl, 0.3 mM TCEP. Two primary peaks were obtained from each sample with major peak at approximately 45kD MW and a minor peak at approximately 79kD. The fractions corresponding to the major peak were pooled and concentrated with an Amicon Ultracel-10 centrifugal filter device (Millipore) and dialyzed vs. 5 mM HEPES, pH 7.5, 50 mM NaCl, 0.3 mM TCEP. Samples collected after dialysis were divided into small aliquots and flash frozen in PCR tubes with liquid nitrogen.

#### Crystallization screening and optimization

Crystallization screening and optimization was conducted in the Collaborative Crystallography Core in the Department of Biochemistry and the University of Wisconsin-Madison. Crystallization experiments were set up using a SPT Labtech mosquito^®^ crystallization robot in MRC SD-2 crystallization plates at 4°C and 20°C (277 and 293 K.) Crystals progressing to diffraction experiments were all obtained at 20°C. Two general screens, Hampton Research IndexHT and Molecular Dimensions JCSG+ were used in this study^46^. Crystals were detected using brightfield and UV fluorescence imaging with a JANSi UVEX-P crystallization plate imaging system supplementing visual inspection with stereomicroscopes. Initial rounds of crystallization optimization were performed in SD2 plates using the mosquito to expand 24 solution conditions by setting columns of experiments in four different sample to reservoir volume ratios. Cryoprotected crystals were harvested in Mitegen micro mounts and flash cooled by immersion in liquid nitrogen.

#### Crystallography

Crystals were screened and X-ray diffraction data was collected at Advanced Photon Source (APS) beamlines LS-CAT and GM/CA@APS, universally on crystals cooled to 100K. Diffraction data was reduced using XDS and scaled with XSCALE^47,48^. Structures were solved by molecular replacement with Phaser within the Phenix suite of programs, automatically rebuilt with phenix.autobuild, iteratively improved with alternating rounds of rebuilding in Coot and refinement using phenix.refine, and validated using MOLPROBITY^49–53^.

**7K1A** crystals providing diffraction data were grown by mixing 200 nL of protein at 9.7 mg/mL in sample buffer (5mM HEPES pH 7.5, 50 mM NaCl, 0.3 mM TCEP) with 150 nL of reservoir solution, was equilibrated against 150 nL 20% MEPEG, 0.2M MgCl2, 0.1M bistris HCL pH 6.5 equilibrated against 50 microliters of reservoir solution in a SD2 plate. Samples were cryoprotected with reservoir solution supplemented to 35% MEPEG 2000. A 360° sweep of data (720 frames) was collected on a MAR 300 CCD detector at LS-CAT beamline 21ID-G on 2018-12-16 using 0.97856 Å X-rays. The phase problem was solved using 2UXU(A) as a molecular replacement model^27^.

**7K1C** crystals of wild-type TtgR with resveratrol were prepared by incubating 0.41 mM protein (9.8 mg/mL) and 0.5 mM resveratrol dissolved in sample buffer for 30 minutes at room temperature prior to setting up crystallization experiments. The crystal yielding the best diffraction data was grown by mixing 200 nL of the protein-ligand sample with 250 nL reservoir (18% PEG4000, 0.2M MgCl2, 0.1M bistris HCl pH 6.5) equilibrated against 50 microliters of reservoir a SD2 plate. Samples were cryoprotected with reservoir solution supplemented with 35% PEG4000. A 360° sweep of data (720 frames) was collected on a MAR 300 CCD detector at LS-CAT beamline 21ID-G on 2018-12-16 using 0.97856 Å X-rays. The phase problem was solved using 2XDN as a molecular replacement model.

**7KD8** crystals were prepared by incubating 0.43 mM (10.4 mg/mL) quadruple mutant protein with 1 mM resveratrol in sample buffer for 30 minutes prior to setting up crystallization experiments. Crystals providing the reported diffraction data set grew from 2 microliters of sample mixed with 2 microliters of reservoir solution (12% MEPEG 2000, 5% 2-methyl-2,4-pentanediol, 0.3 M MgCl2, 0.1 M bistris buffer at pH 6.5 equilibrated in a hanging drop experiment using a siliconized glass cover slip. Samples were cryoprotected with reservoir solution supplemented to 30% MEPEG 2000. A 360°(3600 frames) shutterless data set was collected at LS-CAT 21ID-D on 2019-05-30 with an Eiger 9M direct detector and 1.07812 Å X-rays. The phase problem was solved using 7K1A as a molecular replacement model.

#### Python scripts

All figures were generated using the Matplotlib module in Python 2.7^54^. Scripts used in data analysis and figure generation can be found at: https://github.com/raman-lab/epistasis

---

**Supplementary Table 1:** Control additive data set. A set of random values that are additive with respect to the mean. The standard deviation of each datapoint is 30% of the mean. This control set was used to calculate the R^2^ for comparison to the subnetwork Bahadur expansions.

| Binary | Average | St. Dev |
| --- | --- | --- |
| 00 | 3.1 | 0.93 |
| 01 | 24.5 | 7.35 |
| 10 | 56.1 | 16.83 |
| 11 | 74.4 | 22.32 |

**Supplementary Table 2:**
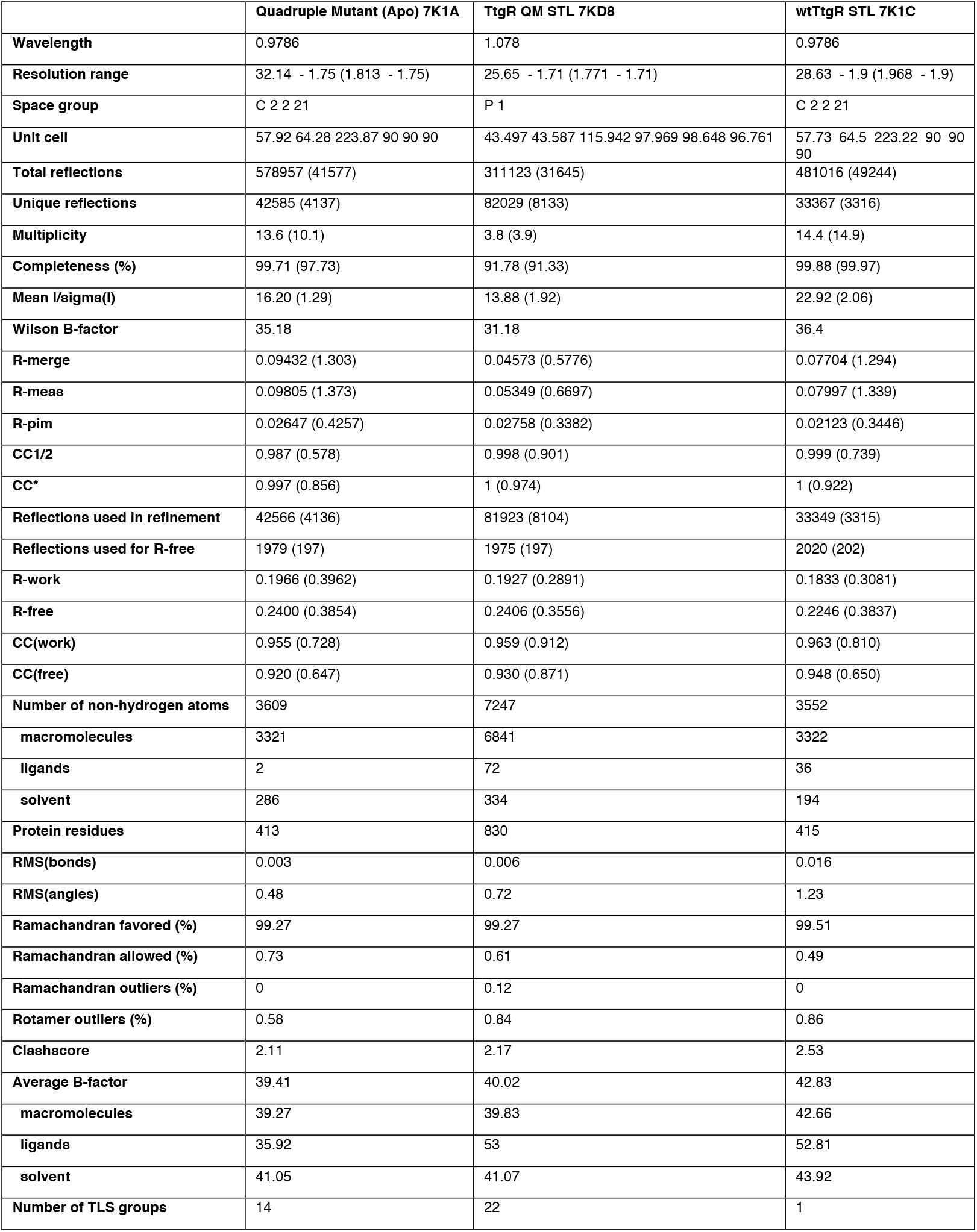
Crystallography refinement statistics. Refinement statistics for three structures: apo quadruple mutant TtgR (7K1A), wildtype TtgR bound to resveratrol (7K1C), and quadruple mutant TtgR bound to resveratrol (7KD8). Statistics for the highest resolution shell are shown in parentheses.

|  | Quadruple Mutant (Apo) 7K1A | TtgR QM STL 7KD8 | wtTtgR STL 7K1C |
| --- | --- | --- | --- |
| Wavelength | 0.9786 | 1.078 | 0.9786 |
| Resolution range | 32.14 - 1.75 (1.813 - 1.75) | 25.65 - 1.71 (1.771 - 1.71) | 28.63 - 1.9 (1.968 - 1.9) |
| Space group | C 2 2 21 | P 1 | C 2 2 21 |
| Unit cell | 57.92 64.28 223.87 90 90 90 | 43.497 43.587 115.942 97.969 98.648 96.761 | 57.73 64.5 223.22 90 90 90 |
| Total reflections | 578957 (41577) | 311123 (31645) | 481016 (49244) |
| Unique reflections | 42585 (4137) | 82029 (8133) | 33367 (3316) |
| Multiplicity | 13.6 (10.1) | 3.8 (3.9) | 14.4 (14.9) |
| Completeness (%) | 99.71 (97.73) | 91.78 (91.33) | 99.88 (99.97) |
| Mean I/sigma(I) | 16.20 (1.29) | 13.88 (1.92) | 22.92 (2.06) |
| Wilson B-factor | 35.18 | 31.18 | 36.4 |
| R-merge | 0.09432 (1.303) | 0.04573 (0.5776) | 0.07704 (1.294) |
| R-meas | 0.09805 (1.373) | 0.05349 (0.6697) | 0.07997 (1.339) |
| R-pim | 0.02647 (0.4257) | 0.02758 (0.3382) | 0.02123 (0.3446) |
| CC1/2 | 0.987 (0.578) | 0.998 (0.901) | 0.999 (0.739) |
| CC* | 0.997 (0.856) | 1 (0.974) | 1 (0.922) |
| Reflections used in refinement | 42566 (4136) | 81923 (8104) | 33349 (3315) |
| Reflections used for R-free | 1979 (197) | 1975 (197) | 2020 (202) |
| R-work | 0.1966 (0.3962) | 0.1927 (0.2891) | 0.1833 (0.3081) |
| R-free | 0.2400 (0.3854) | 0.2406 (0.3556) | 0.2246 (0.3837) |
| CC(work) | 0.955 (0.728) | 0.959 (0.912) | 0.963 (0.810) |
| CC(free) | 0.920 (0.647) | 0.930 (0.871) | 0.948 (0.650) |
| Number of non-hydrogen atoms | 3609 | 7247 | 3552 |
| macromolecules | 3321 | 6841 | 3322 |
| ligands | 2 | 72 | 36 |
| solvent | 286 | 334 | 194 |
| Protein residues | 413 | 830 | 415 |
| RMS(bonds) | 0.003 | 0.006 | 0.016 |
| RMS(angles) | 0.48 | 0.72 | 1.23 |
| Ramachandran favored (%) | 99.27 | 99.27 | 99.51 |
| Ramachandran allowed (%) | 0.73 | 0.61 | 0.49 |
| Ramachandran outliers (%) | 0 | 0.12 | 0 |
| Rotamer outliers (%) | 0.58 | 0.84 | 0.86 |
| Clashscore | 2.11 | 2.17 | 2.53 |
| Average B-factor | 39.41 | 40.02 | 42.83 |
| macromolecules | 39.27 | 39.83 | 42.66 |
| ligands | 35.92 | 53 | 52.81 |
| solvent | 41.05 | 41.07 | 43.92 |
| Number of TLS groups | 14 | 22 | 1 |

**Supplementary Table 3:** Psi values for Bahadur Expansion. Psi values for all orders of interactions (right column) for each mutant.

|  | 0000 | 0001 | 0010 | 0011 | 0100 | 0101 | 0110 | 0111 | 1000 | 1001 | 1010 | 1011 | 1100 | 1101 | 1110 | 1111 | Residue Interactions |
| --- | --- | --- | --- | --- | --- | --- | --- | --- | --- | --- | --- | --- | --- | --- | --- | --- | --- |
| $\Psi_0$ | 1 | 1 | 1 | 1 | 1 | 1 | 1 | 1 | 1 | 1 | 1 | 1 | 1 | 1 | 1 | 1 | 0 |
| $\Psi_1$ | -1 | -1 | -1 | -1 | -1 | -1 | -1 | -1 | 1 | 1 | 1 | 1 | 1 | 1 | 1 | 1 | 1 |
| $\Psi_2$ | -1 | -1 | -1 | -1 | 1 | 1 | 1 | 1 | -1 | -1 | -1 | -1 | 1 | 1 | 1 | 1 | 2 |
| $\Psi_3$ | -1 | -1 | 1 | 1 | -1 | -1 | 1 | 1 | -1 | -1 | 1 | 1 | -1 | -1 | 1 | 1 | 3 |
| $\Psi_4$ | -1 | 1 | -1 | 1 | -1 | 1 | -1 | 1 | -1 | 1 | -1 | 1 | -1 | 1 | -1 | 1 | 4 |
| $\Psi_5$ | 1 | 1 | 1 | 1 | -1 | -1 | -1 | -1 | -1 | -1 | -1 | -1 | 1 | 1 | 1 | 1 | 1-2 |
| $\Psi_6$ | 1 | 1 | -1 | -1 | 1 | 1 | -1 | -1 | -1 | -1 | 1 | 1 | -1 | -1 | 1 | 1 | 1-3 |
| $\Psi_7$ | 1 | -1 | 1 | -1 | 1 | -1 | 1 | -1 | -1 | 1 | -1 | 1 | -1 | 1 | -1 | 1 | 1-4 |
| $\Psi_8$ | 1 | 1 | -1 | -1 | -1 | -1 | 1 | 1 | 1 | 1 | -1 | -1 | -1 | -1 | 1 | 1 | 2-3 |
| $\Psi_9$ | 1 | -1 | 1 | -1 | -1 | 1 | -1 | 1 | 1 | -1 | 1 | -1 | -1 | 1 | -1 | 1 | 2-4 |
| $\Psi_{10}$ | 1 | -1 | -1 | 1 | 1 | -1 | -1 | 1 | 1 | -1 | -1 | 1 | 1 | -1 | -1 | 1 | 3-4 |
| $\Psi_{11}$ | -1 | -1 | 1 | 1 | 1 | 1 | -1 | -1 | 1 | 1 | -1 | -1 | -1 | -1 | 1 | 1 | 1-2-3 |
| $\Psi_{12}$ | -1 | 1 | -1 | 1 | 1 | -1 | 1 | -1 | 1 | -1 | 1 | -1 | -1 | 1 | -1 | 1 | 1-2-4 |
| $\Psi_{13}$ | -1 | 1 | 1 | -1 | -1 | 1 | 1 | -1 | 1 | -1 | -1 | 1 | 1 | -1 | -1 | 1 | 1-3-4 |
| $\Psi_{14}$ | -1 | 1 | 1 | -1 | 1 | -1 | -1 | 1 | -1 | 1 | 1 | -1 | 1 | -1 | -1 | 1 | 2-3-4 |
| $\Psi_{15}$ | 1 | -1 | -1 | 1 | -1 | 1 | 1 | -1 | -1 | 1 | 1 | -1 | 1 | -1 | -1 | 1 | 1-2-3-4 |

