## Supplementary Figures for "Epistasis shapes the fitness landscape of an allosteric specificity switch"

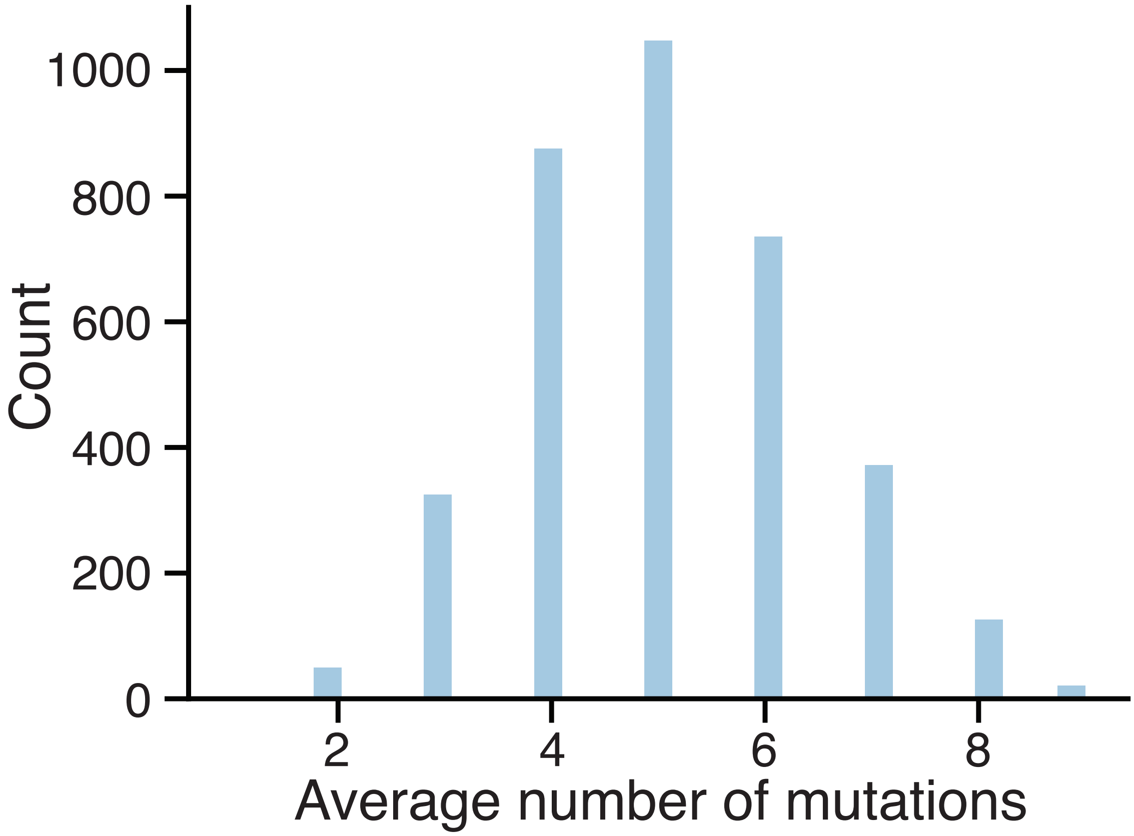


**Supplementary Figure 1: Mutation distribution for synthesized resveratrol designs**
Histogram of the number of mutations in the set of Rosetta designs selected for oligo synthesis. The average number of mutations was 5.07 with a variance of 1.78.

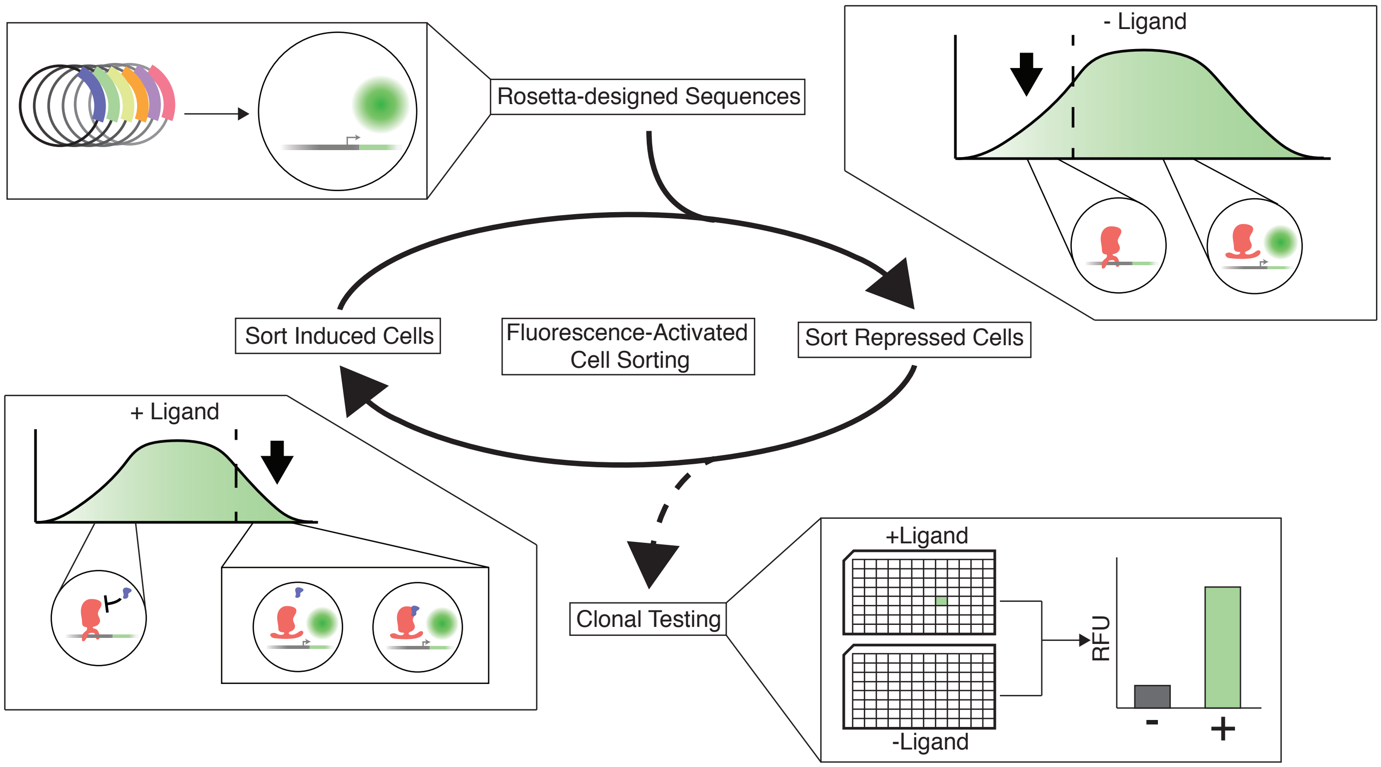


**Supplementary Figure 2: Workflow for screening ligand-specific TtgR variants by fluorescence activated cell sorting**

Rosetta-designed TtgR variants are transformed into E. coli cells carrying the reporter. TtgR variants are sorted by toggling between repressed and induced states (solid arrow). The lower 50% of fluorescent cells are sorted in the absence of inducer to isolate variants that are able to repress transcription. Subsequently, the sorted population are grown and induced with resveratrol. The top 5% of fluorescent cells are sorted to isolate variants capable of binding to the ligand and inducing GFP expression. After toggling multiple times, the repressed sort is repeated a final time before the subpopulation is clonally tested with both naringenin and resveratrol (dashed arrow).

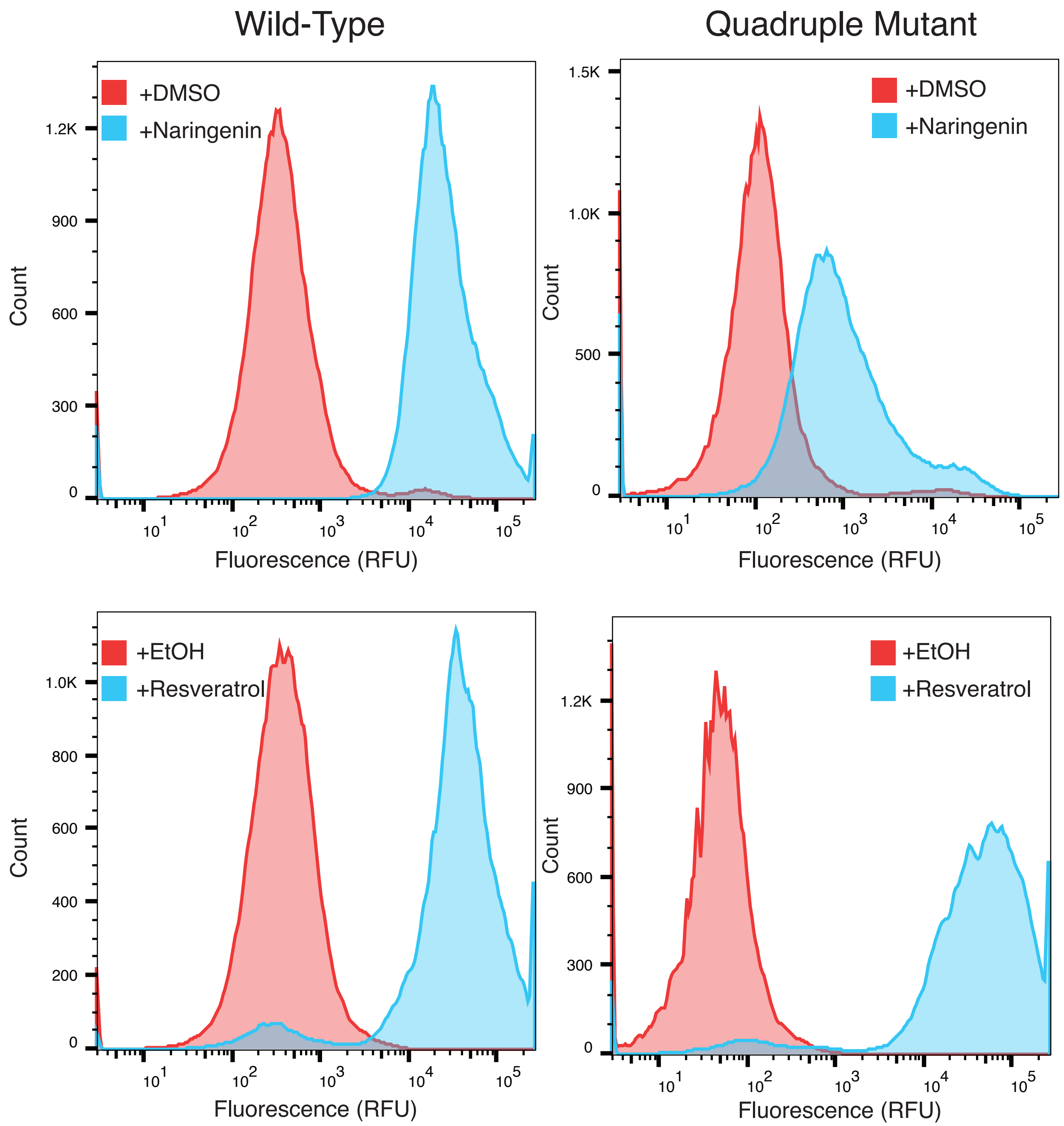


**Supplementary Figure 3: Fluorescence distributions of wildtype TtgR and quadruple mutant**
Flow cytometry histograms of wildtype TtgR and quadruple mutant TtgR with and without inducers. Naringenin 2000μM (blue) dissolved in DMSO and DMSO-only control (red), Resveratrol 250μM (blue) dissolved in ethanol and ethanol-only control (red) are shown.

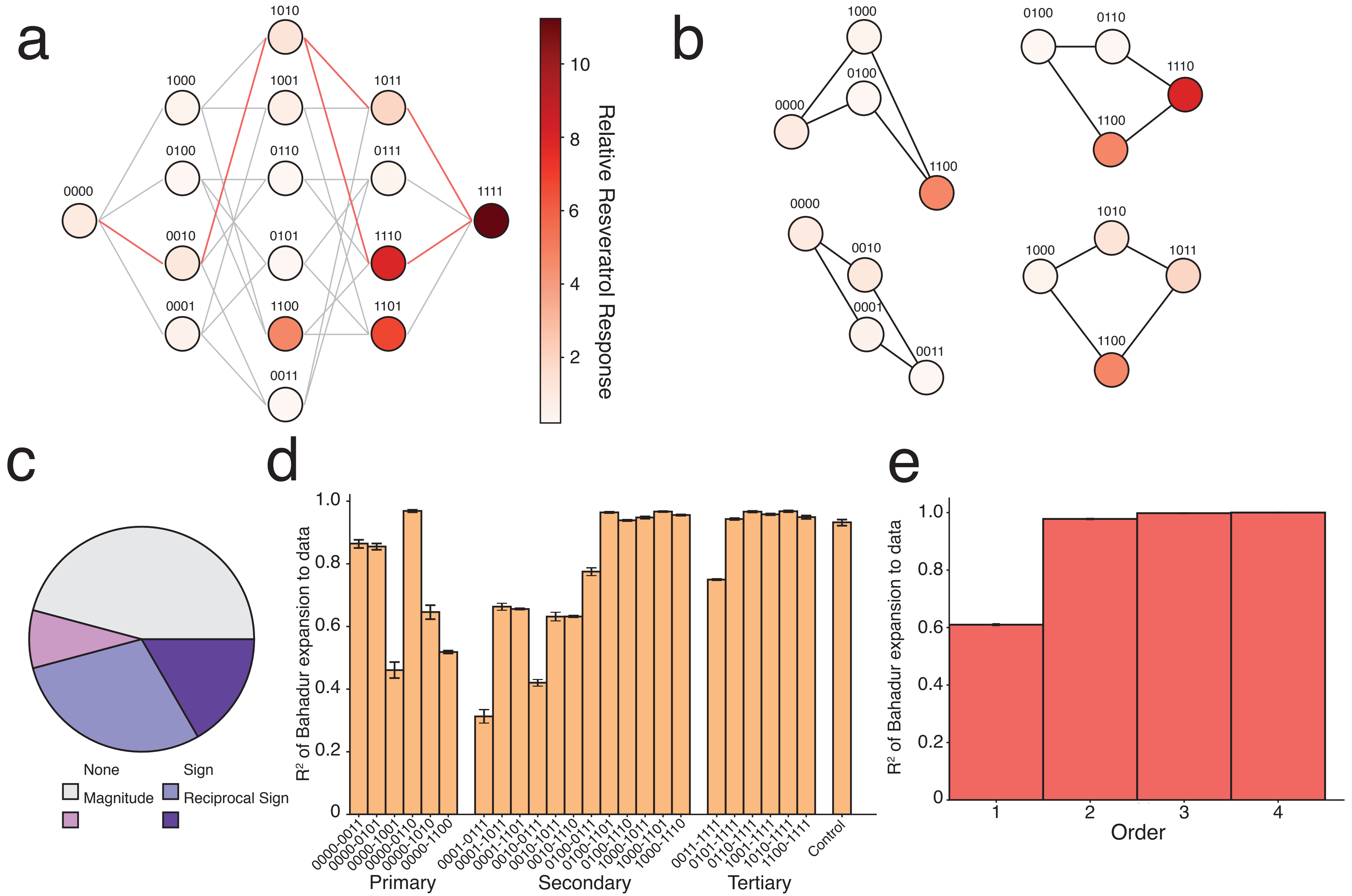


**Supplementary Figure 4: Fitness landscape and epistatic effects of response to resveratrol (No logarithm transformation)**
**(a)** Fitness landscape of resveratrol response for all 16 TtgR variants connecting the wildtype to the quadruple mutant with each variant shown as a node in the graph. Layout and subfigures are analogous to those in Fig. 2. Nodes are shaded by fold induction ratio at 250μM resveratrol of that variant normalized to fold induction ratio of wildtype TtgR. **(b)** Select subnetworks highlighting key epistatic interactions **(c)** Different types of epistasis in the 24 subnetworks. **(d)** R2 values for each subnetwork individually. Subnetworks are labeled based on the first and last node and are separated into primary, secondary, or tertiary groups. The control group is based off of simulated additive data with standard deviations equivalent to 30% of the mean for each node. **(e)** Statistical analysis of the order of epistasis over the entire fitness landscape based on R2 values from the Bahadur expansion. The x-axis indicates the highest order terms included in the Bahadur expansion (Eg: Order 2 contains 1st and 2nd order terms). Error for (d) and (e) are derived from bootstrap confidence intervals (See methods).

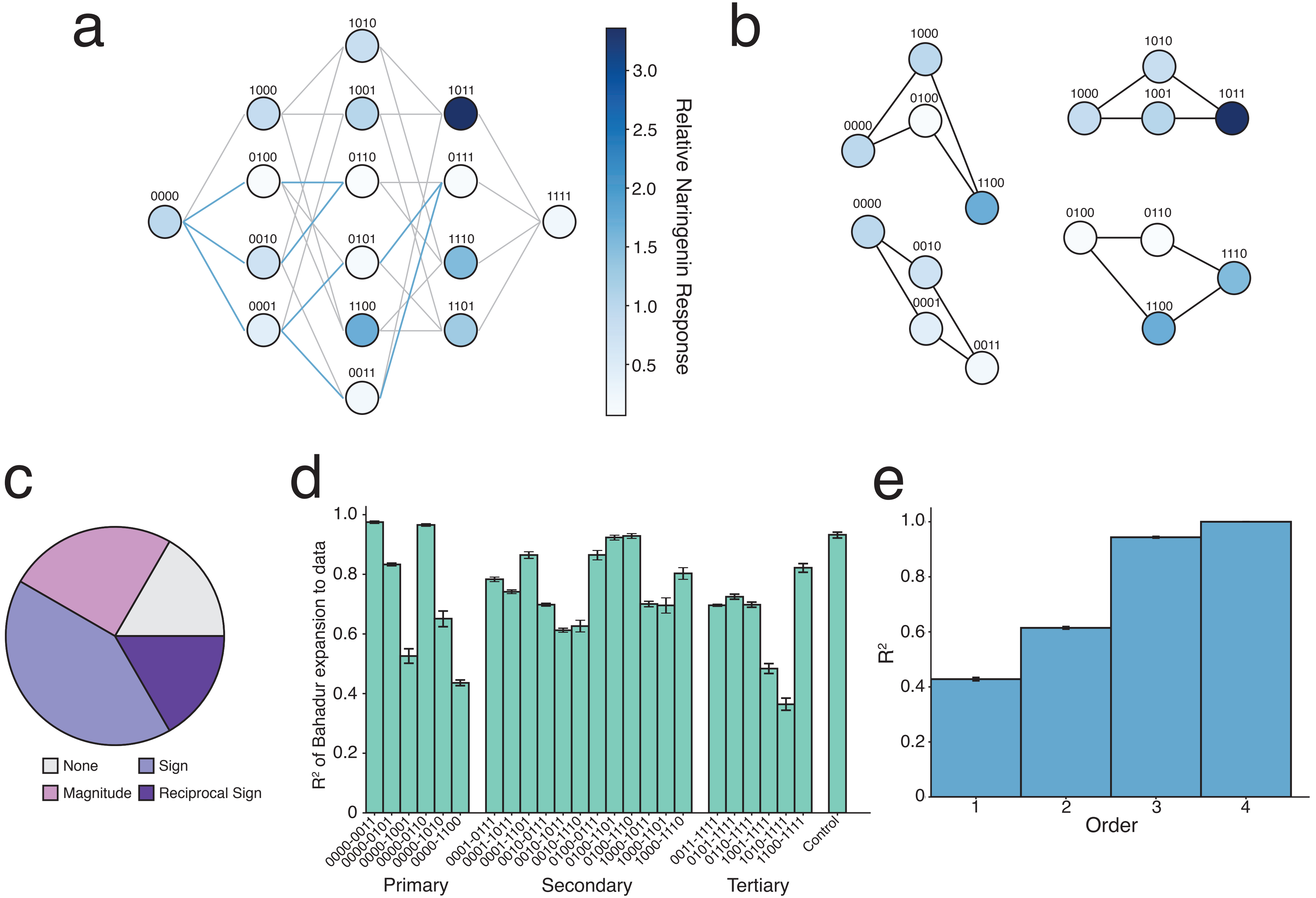


**Supplementary Figure 5: Fitness landscape and epistatic effects of response to naringenin (No logarithm transformation)**

**(a)** Fitness landscape of naringenin response for all 16 TtgR variants connecting wildtype to quadruple mutant with each variant shown as a node in the graph. Labeling and layout are identical to the resveratrol landscape. Nodes separated by a single mutation are connected by edges showing viable (bold blue) and unviable paths (light gray) through sequence space. Nodes are shaded by fold induction ratio at 2000μM naringenin of that variant normalized to fold induction ratio of wildtype TtgR. **(b)** Select subnetworks highlighting key epistatic interactions **(c)** Different types of epistasis in the 24 subnetworks. **(d)** R2 values for each subnetwork individually quantified using the Bahadur expansion. Labels, grouping, and the control set are identical to 3d. **(e)** Statistical analysis of the order of epistasis over the entire fitness landscape based on R2 values from the Bahadur expansion. Labeling is identical to 3e.

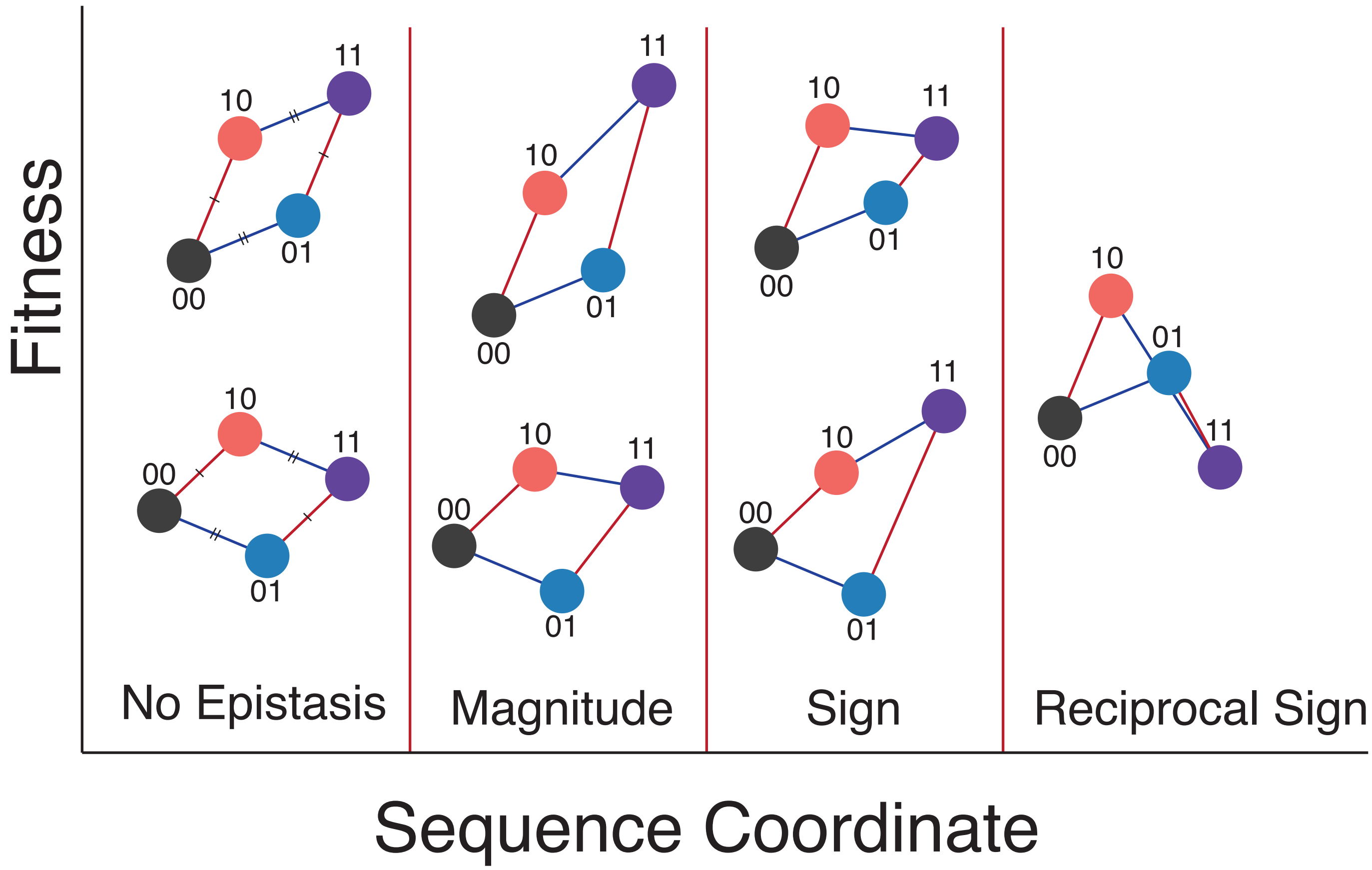


**Supplementary Figure 6: Visual definition of different types of epistasis**

Visual representation of different types of epistasis. This graphical representation separates example subnetworks based on the type of epistasis. An arbitrary fitness metric is plotted against a sequence coordinate where each mutation is represented by a binary string. A system is non-epistatic when the combined effect of mutations is the sum of their individual effects. Magnitude epistasis occurs when the combined effect of mutations is greater than the sum of their individual effects (no change in direction). Sign epistasis occurs when one mutation switches direction from beneficial to detrimental (or vice versa) depending on the background in which it is introduced. Reciprocal sign epistasis occurs when both mutations switch direction depending on the background.

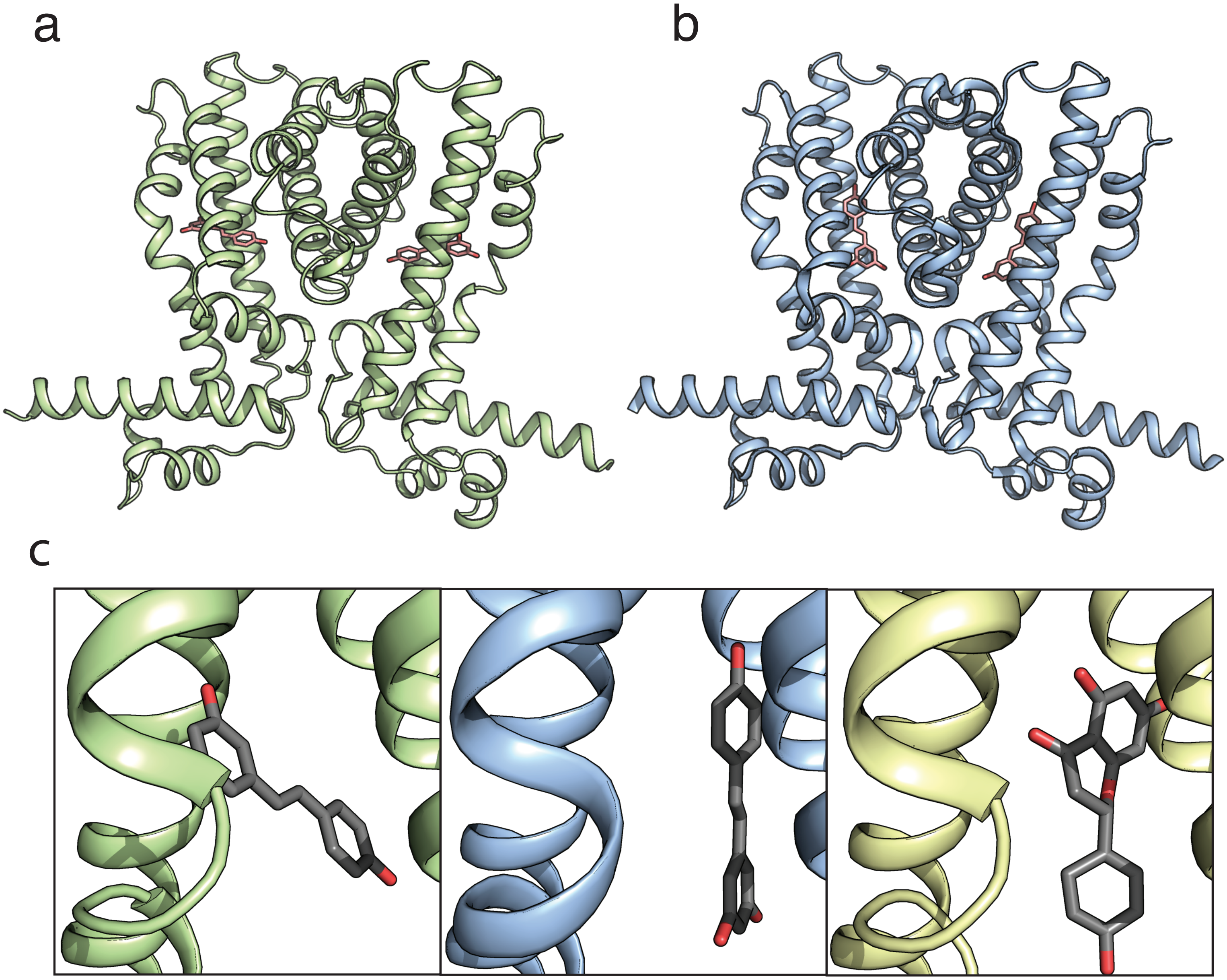

**Supplementary Figure 7: Whole structure of ligand-bound wildtype and quadruple mutant**

TtgR is as an all-helical dimer. The helix-turn-helix domain at the N-terminal end binds to DNA. The ligand binding pocket is enclosed by five angled helices. An additional helix at the C-terminal end forms the dimerization interface. The quadruple mutant (PDB: 7KD8) **(a)** is structurally identical to the wildtype (PDB: 7K1C) **(b)**. Resveratrol is shown as pink sticks in both. **(c)** A close-up view of the binding orientation of resveratrol in the pocket. The quadruple mutant (left) binds to resveratrol in the horizontal orientation. Wildtype TtgR binds to resveratrol (middle) or to naringenin (right, PDB ID: 2UXU) in the vertical orientation.

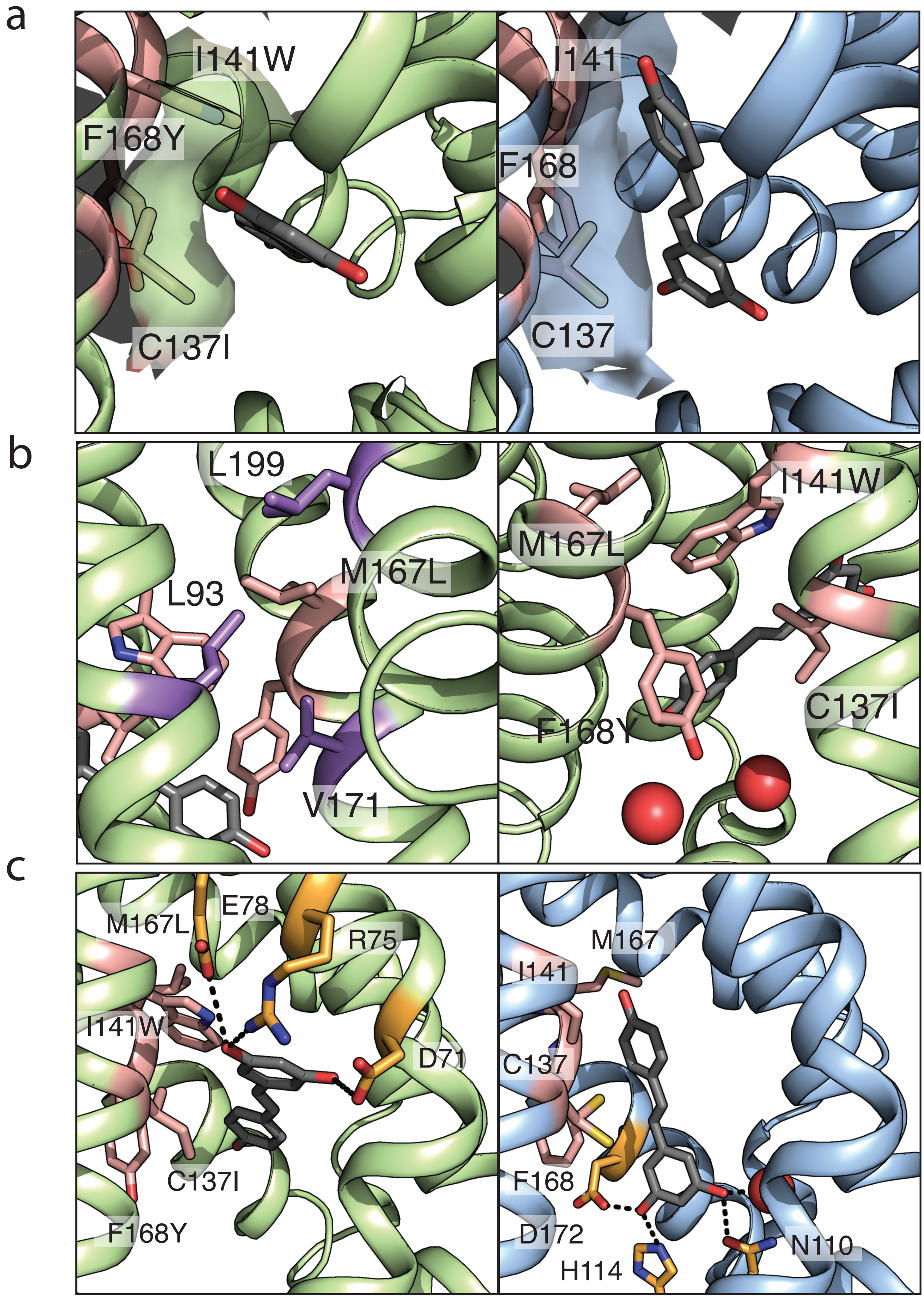


**Supplementary Figure 8: Interactions of mutated positions and alternate hydrogen bonding networks**
**(a)** The C137I mutation creates a small cleft in the binding pocket that can enhance shape complementarity to resveratrol in the horizontal binding mode. The quadruple mutant (left) is shown in comparison to wildtype (right). The van der Waals surface of residue 137 and 141 is shown for both structures. **(b)** (Left) M167L creates nonpolar interactions with residues in helices composing the binding pocket and dimerization interface. These residues are labeled and shown as purple sticks. Mutated positions 137, 141, 167, and 168 are shown in red. 167 also plays a role in positioning the I141W side chain. (Right) The F168Y substitution enables the formation of additional hydrogen bonds to solvent that can create a hydrogen bond network with D172. **(c)** The hydrogen bond network differs for the quadruple mutant between chain A and chain B due to the slightly different position of the resveratrol molecules in each. (Left) In the quadruple mutant, D71, R75 and E78 make hydrogen bonds with the resveratrol molecules. (Right) The hydrogen bonding network of wildtype chain A is identical to chain B.

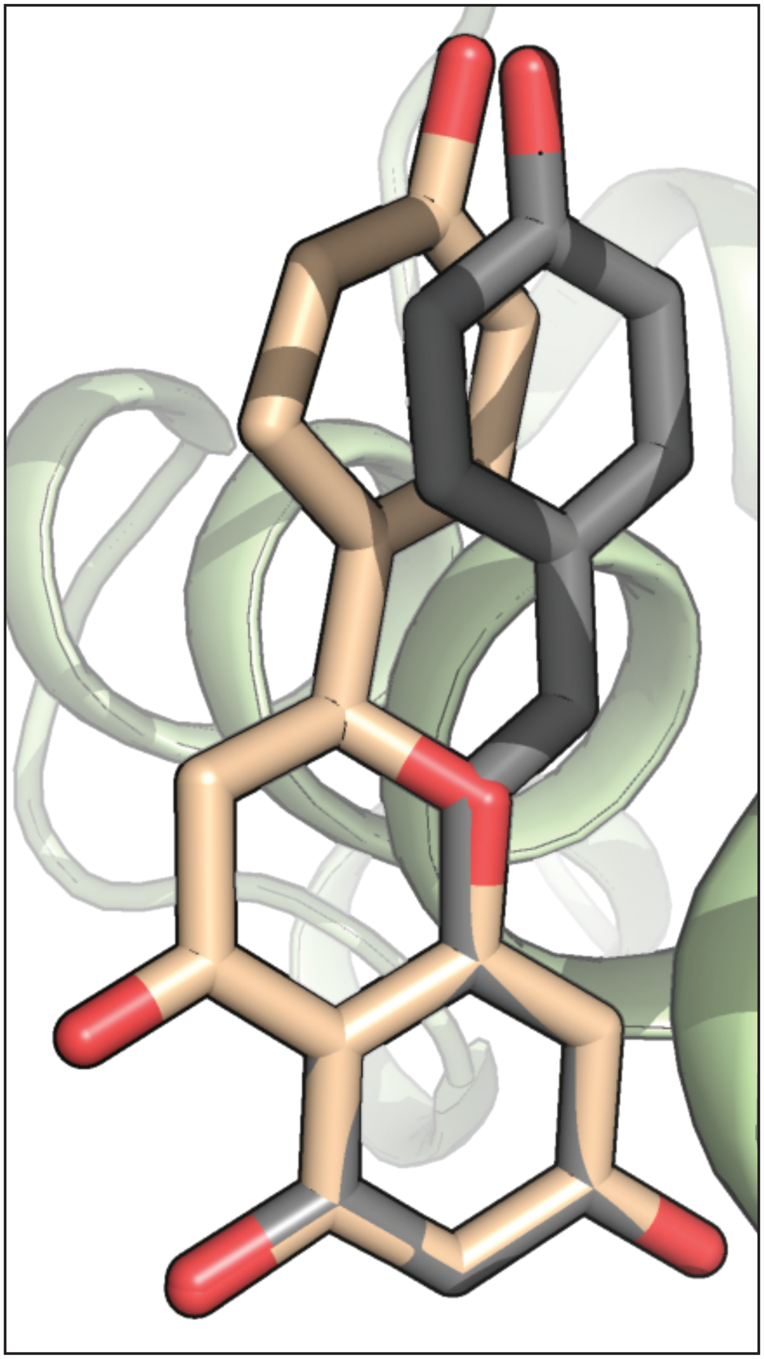

**Supplementary Figure 9: Naringenin and resveratrol overlap**
Naringenin (in brown) derived from a previous structure (PDB: 2UXU) is overlapped with resveratrol (in grey) via the pair_fit function in Pymol. The structures of each ligand are similar with respect to the location of hydroxyl groups, but differ by the addition of a carbonyl in the 4-chromanone backbone of naringenin.

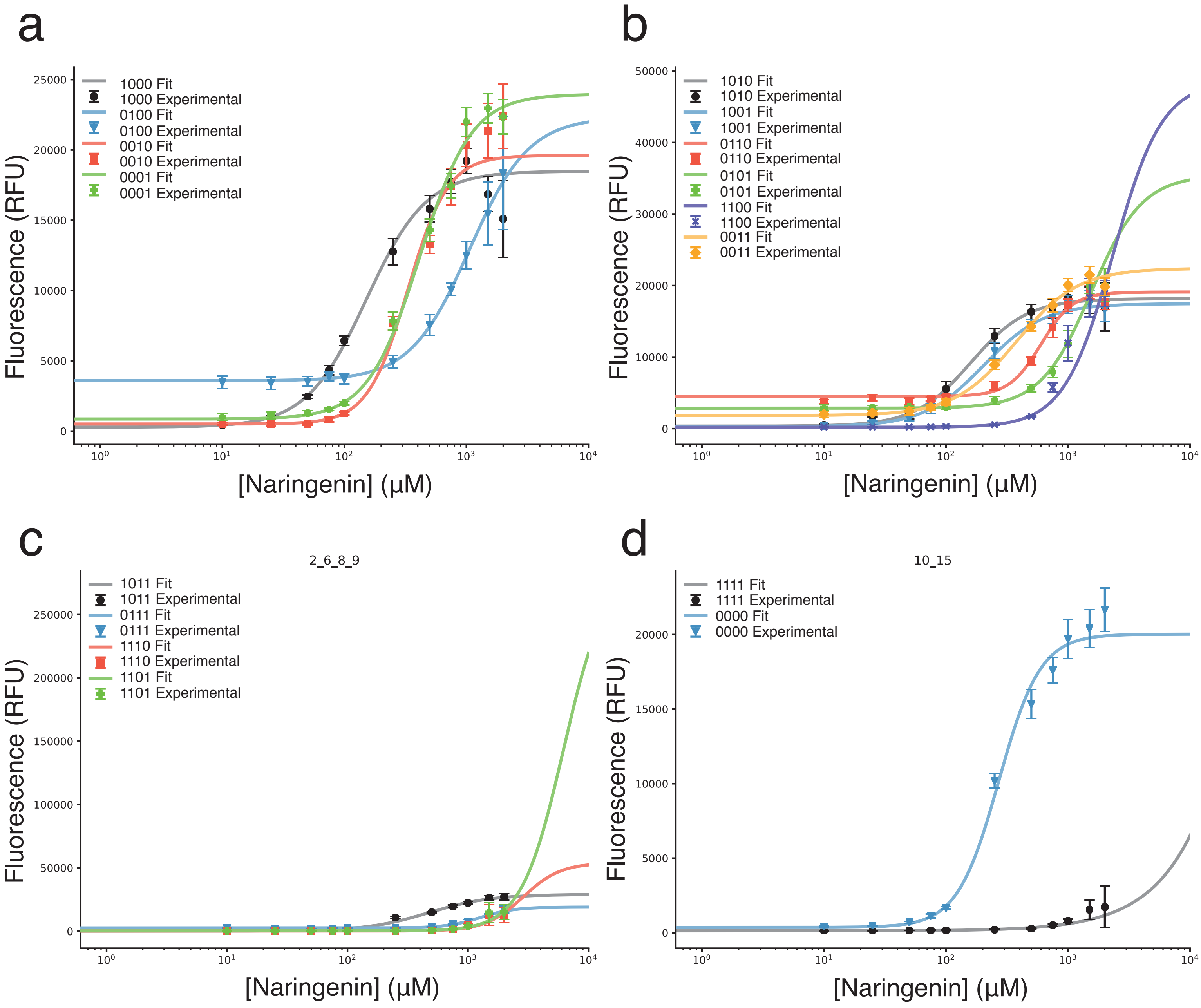


**Supplementary FIgure 10: Naringenin dose response curves**
Dose response curves to naringenin for all 16 mutational combinations. Naringenin concentration varied between 0μM and 2000μM naringenin. Averages and standard deviations of triplicate measurements are shown unless otherwise specified (see Methods). **(a)** Single mutant fits (1000, 0100, 0010, and 0001). Fit is shown as a solid line and experimental data is shown as markers with error bars. **(b)** Double mutant fits (1010, 1001, 0110, 1100, and 0011). **(c)** Triple mutant fits (1011, 0111, 1110, and 1101). **(d)** Wildtype (0000) and quadruple mutant (1111) dose response curves.

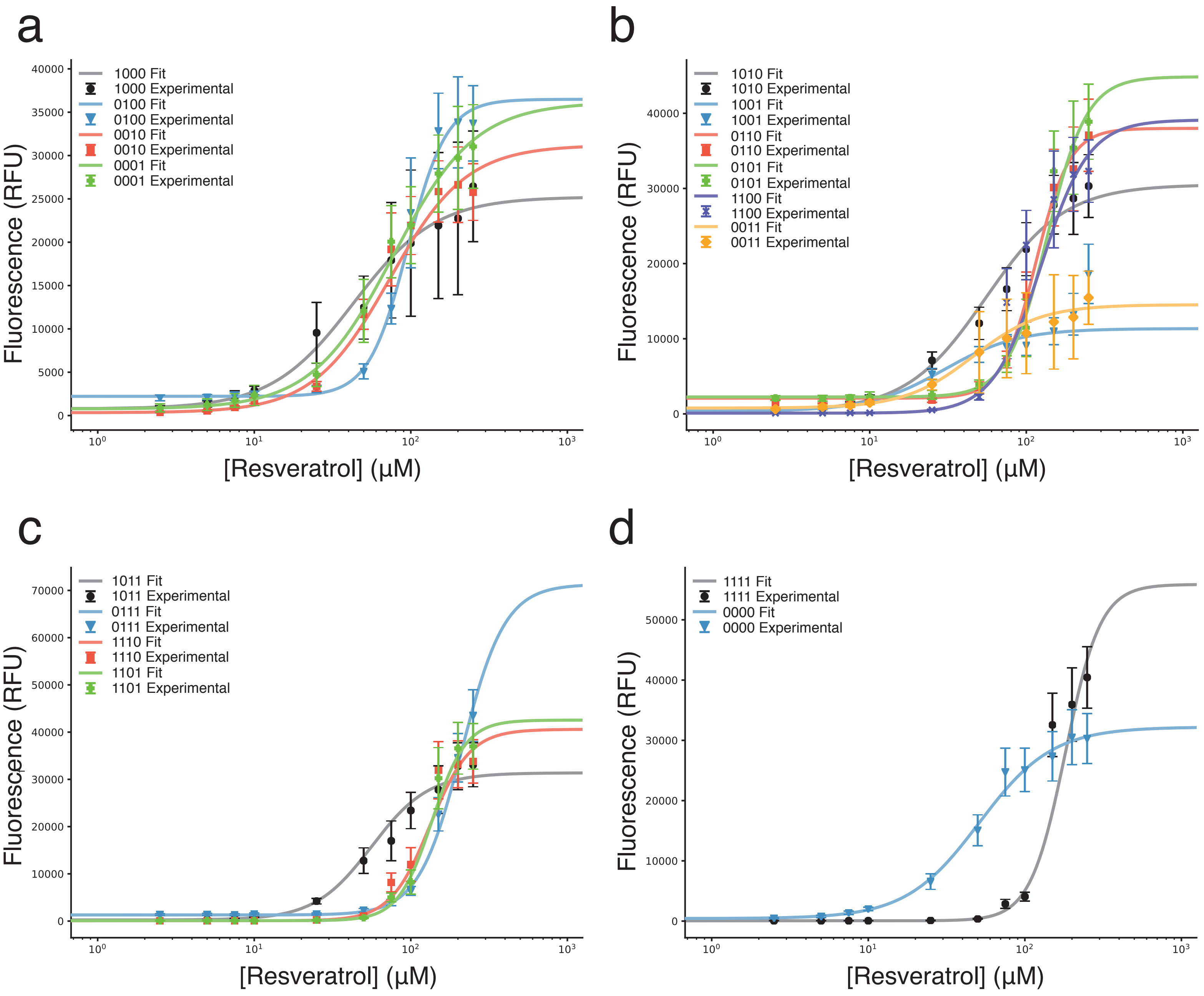


**Supplementary Figure 11: Resveratrol dose response curves**
Dose response curves for all 16 mutational combinations to resveratrol. Resveratrol concentration varied between 0μM and 250μM resveratrol. Averages and standard deviations of triplicate measurements are shown unless otherwise specified (see Methods). **(a)** Single mutant fits (1000, 0100, 0010, and 0001). Fit is shown as a solid line and experimental data is shown as markers with error bars. **(b)** Double mutant fits (1010, 1001, 0110, 1100, and 0011). **(c)** Triple mutant fits (1011, 0111, 1110, and 1101). **(d)** Wildtype (0000) and quadruple mutant (1111) dose response curves.

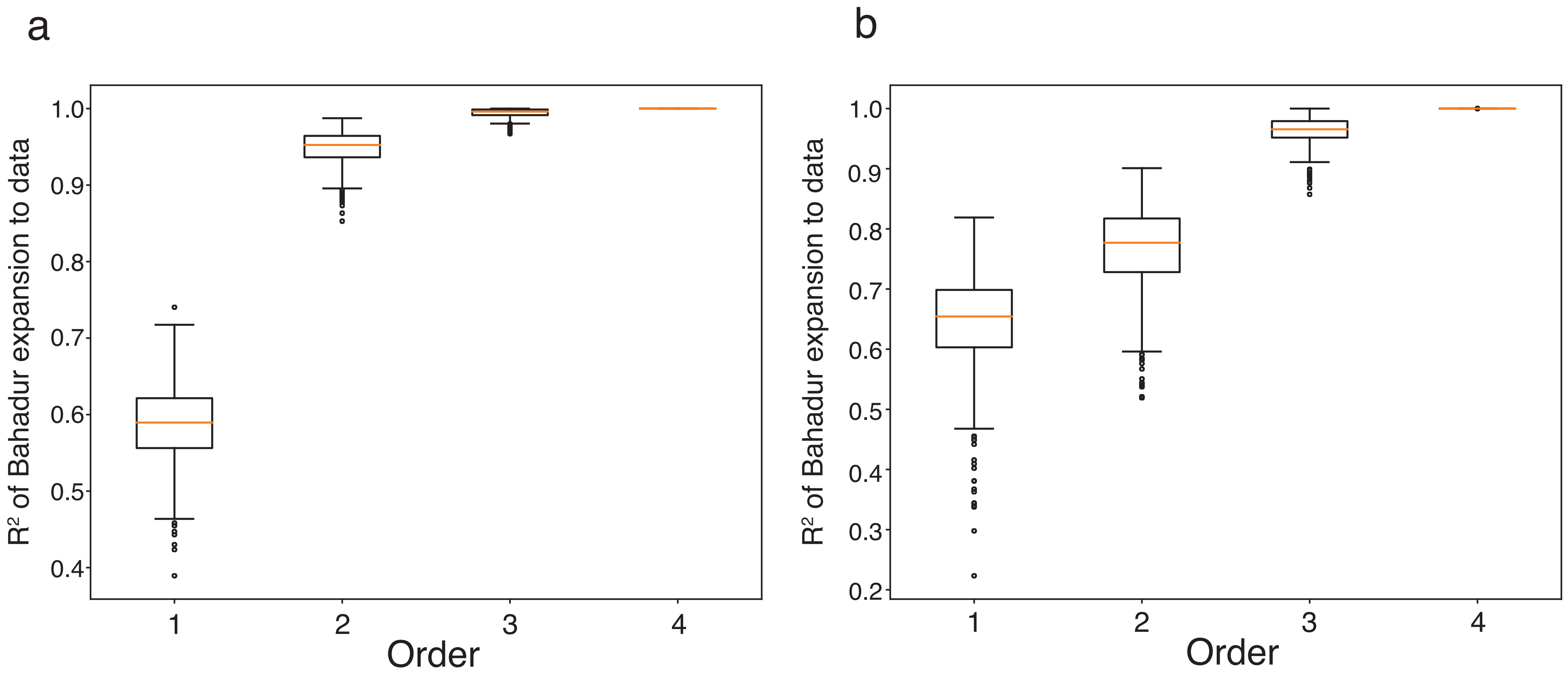


**Supplementary Figure 12: Distribution of $R^2$ for first and higher order interactions for the full network**
Distribution of $R^2$ values from the Bahadur expansion model applied to the full 16-member network after stochastic sampling (N=500) of $\log_{10}$ fluorescence values based on experimental averages and standard deviations. Boxplots of first, second, third, and fourth order interactions are shown for **(a)** resveratrol and **(b)** naringenin. The orange line is the median $R^2$ value for the distribution and the box encloses the interquartile range (IQR). The whiskers extend to the $R^2$ value closest outside 1.5 times the IQR. Circles represent data points outside the whiskers.

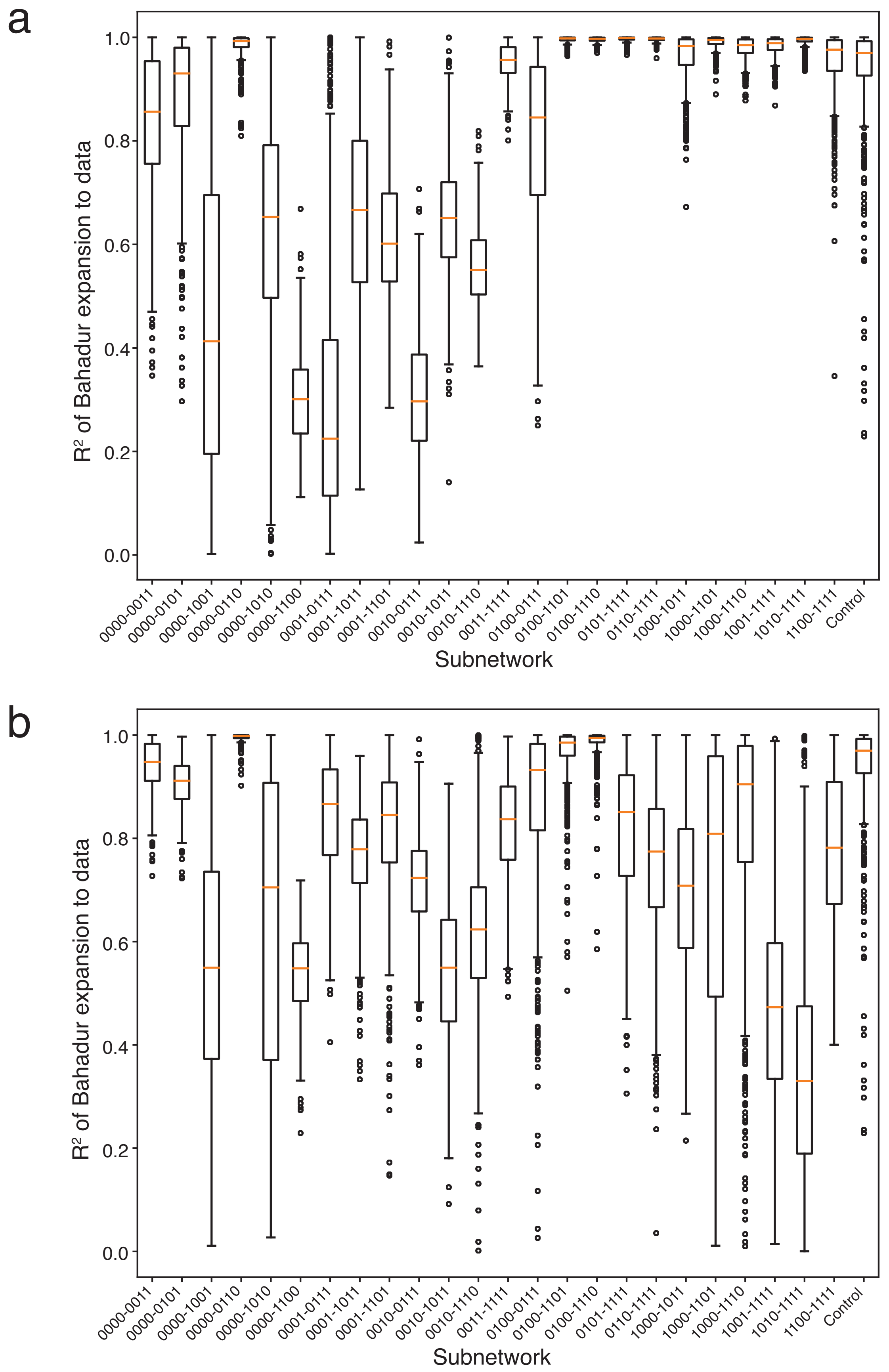


**Supplementary Figure 13: Distribution of $R^2$ for first and second order interactions for individual subnetworks**
Distribution of $R^2$ values from the Bahadur expansion model applied to each subnetwork after stochastic sampling of experimental $\log_{10}$ fluorescence values of response to **(a)** resveratrol or **(b)** naringenin based on experimental averages and standard deviations. Each subnetwork network was modeled 500 times by Monte Carlo sampling. The orange line is the median $R^2$ value for the distribution and the box encloses the interquartile range (IQR). The whiskers extend to the $R^2$ value closest outside a1.5 times the IQR. Circles represent data points outside the whiskers.

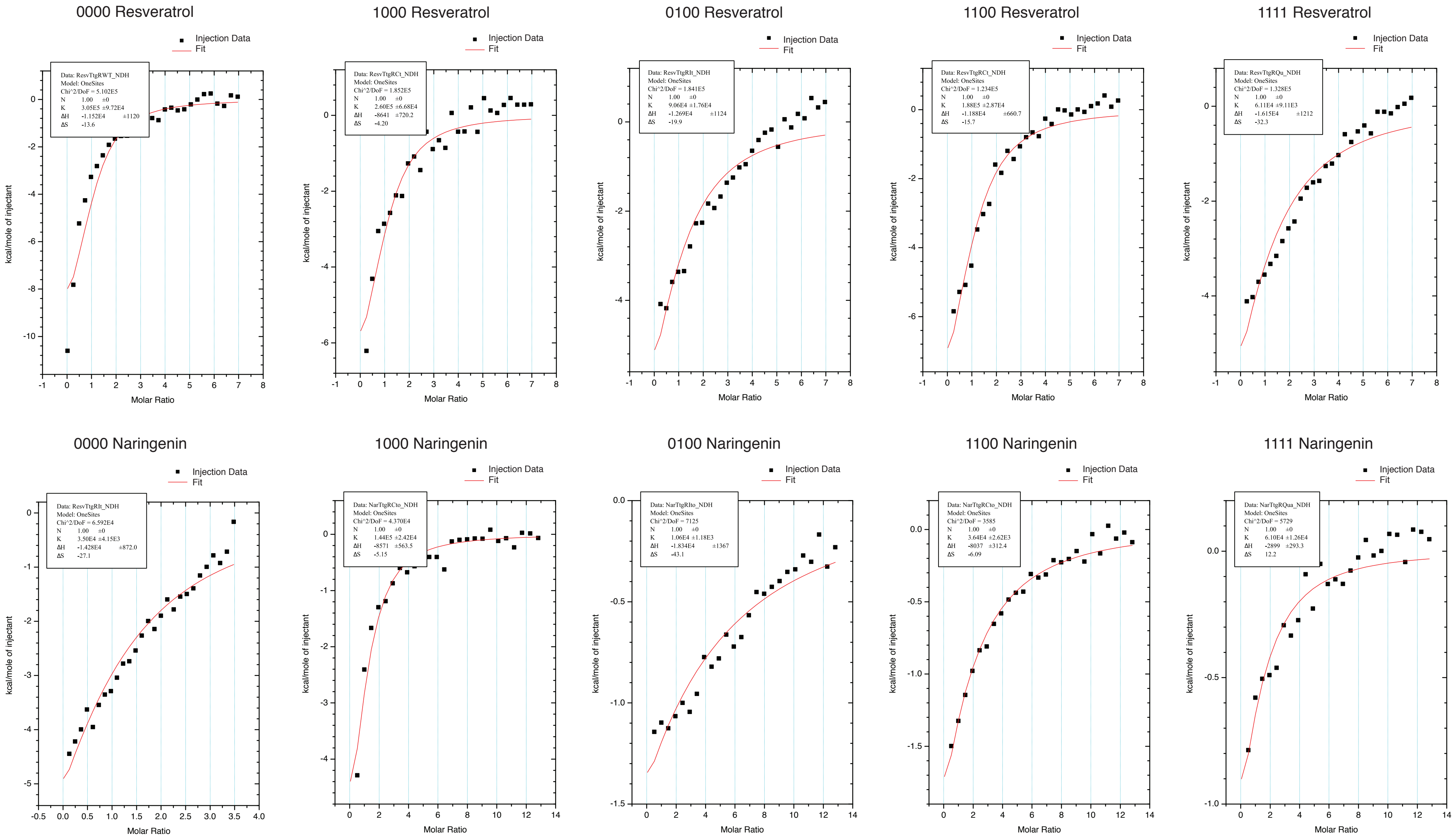


**Supplementary Figure 14: Estimating binding parameters from isothermal calorimetry of wildtype TtgR and variants**
Isothermal titration calorimetry experimental data for affinity of TtgR mutants to either naringenin or resveratrol. Heat per mole of ligand injected (kCal/mol) is plotted as a function of the molar ratio of ligand:protein. Binding parameters are estimated from single site binding model fits using Origin 7.0 software (MicroCal). Due to low affinities for both naringenin and resveratrol, stoichiometry was fixed to 1 for both naringenin and resveratrol (see methods).
